# Fidelity of Spatiotemporal Patterns of Brain Activity Across Sampling Rate, Scan Duration, and Frequency Content

**DOI:** 10.64898/2026.01.02.697199

**Authors:** Theodore J. LaGrow, Harrison Watters, Lauren Daley, Vaibhavi Itkyal, Dolly Seeburger, Nmachi Anumba, Abia Fazili, Michael A. Kelberman, Vince Calhoun, Wen-Ju Pan, Eric H. Schumacher, Shella Keilholz

## Abstract

Intrinsic brain activity is characterized by large-scale spatiotemporal patterns that underpin functional connectivity and cognition. Quasi-periodic patterns (QPPs) and complex principal component analysis (cPCA) have emerged as reproducible methods for capturing spatiotemporal network interactions in resting-state functional magnetic resonance imaging (rs-fMRI). However, these methods remain sensitive to methodological factors such as scan duration, repetition time (TR), and frequency band selection. This study systematically evaluates how these parameters influence the stability and reliability of QPP- and cPCA-derived functional connectivity patterns across multiple datasets. Using five independent rs-fMRI datasets, we evaluate the impact of scan length on pattern reliability, explore the effects of TR on spatiotemporal patterns, and compare the sensitivity of different frequency bands (Slow-5, Slow-4, infraslow) in capturing network dynamics. Our findings reveal that while both QPPs and cPCA detect intrinsic network activity, their reliability varies with acquisition parameters. QPPs exhibit greater stability in shorter scans, making them suitable for individual-level analyses, whereas cPCA provides a broader representation of phase-coherent fluctuations but shows greater between-subject variability and benefits more from longer, group-level acquisitions. Additionally, frequency band selection significantly influences the temporal structure of extracted patterns: in our analyses, Slow-5 (0.01–0.027 Hz) tended to emphasize more recurrent, synchronized network configurations, whereas Slow-4 (0.027–0.073 Hz) more often revealed transitions between connectivity states. These results provide critical insights into optimizing methodological choices for dynamic functional connectivity analysis, enhancing the interpretability of spatiotemporal patterns in both basic and clinical neuroimaging research.

## 1 Introduction

Intrinsic brain activity is composed of large-scale spatiotemporal patterns that coordinate the interaction of functional networks. These patterns reflect coordinated activity across distributed brain regions and play a critical role in cognition, perception, and neurological disorders (Calhoun et al., 2014; S. Keilholz et al., 2017; Ma et al., 2014; Preti et al., 2017). Observed using resting-state fMRI (rs-fMRI), these dynamics provide insight into the brain’s functional architecture Capturing such dynamics requires time-resolved analysis, but no community consensus exists on which method yields the most reliable picture (Iraji et al., 2020, 2021). Sliding-window correlation (SWC) continuously estimates connectivity over shifting windows, yet its performance depends sensitively on window length and frequency content (Cabral et al., 2017; Chang & Glover, 2010; S. D. Keilholz et al., 2013; Preti et al., 2017; Shakil et al., 2016; Yaesoubi et al., 2015).

State-based approaches such as Hidden Markov Models (HMMs), co-activation patterns (CAPs), and clustering algorithms instead identify recurring “brain states,” producing a discrete view of network transitions (Allen et al., 2014; X. Liu et al., 2018; Vidaurre et al., 2017). Two contemporary entrants—quasi-periodic patterns (QPPs) and complex principal component analysis (cPCA)—offer complementary frameworks for extracting structured spatiotemporal motifs. QPPs iteratively isolate a recurrent propagation template (Majeed et al., 2009; Yousefi & Keilholz, 2021; Yousefi et al., 2018), whereas cPCA exploits the complex domain to capture phase-coherent modes (Bolt et al., 2022). Despite distinct mathematics, both often recover strikingly similar large-scale patterns (Bolt et al., 2022). Yet we still lack a head-to-head evaluation of how acquisition and analytic parameters—particularly scan duration, TR, window length, and frequency band selection—shape the patterns each method produces. Addressing how scan length, TR, and frequency band influence QPP- and cPCA-derived patterns across datasets is the central aim of the present study.

Spatiotemporal patterns of brain activity appear remarkably consistent across species, linking fluctuations in rodent cortex and the human brain to underlying infraslow neural activity (Grooms et al., 2017; Thompson et al., 2014). These patterns track canonical resting state networks and the global signal (Belloy et al., 2018), persist during neuromodulatory perturbations such as optogenetic stimulation of the locus coeruleus (Anumba et al., 2024), and even survive the species divide between anesthetized rodents and awake humans (Xu et al., 2022). However, this apparent universality is tempered by an overlooked issue: to extract spatiotemporal patterns with the QPP algorithm is not a single template but a family of solutions whose exact form depends on analyst choices, including window length, frequency band, initialization, preprocessing the way phase information is handled. Shorter windows (e.g., *<*30 s) can sharpen temporal resolution but tend to magnify noise and reduce the stability of estimated templates, whereas substantially longer windows (e.g., *>* 60 s) average over fast propagations and bias the patterns toward more slowly varying components (Shakil et al., 2016). Comparable trade-offs arise when removing slow drifts or when selecting the phase threshold that defines cPCA modes. Without a systematic evaluation, we cannot know how these parameter decisions shape the biological story we tell.

In human studies, the spatiotemporal patterns have been linked to fluctuations in arousal and attentional states, propagating as global waves through the neocortex and thalamus (Raut et al., 2021). Supporting this, QPP-based metrics differentiate control from ADHD groups (Abbas et al., 2019), as well as sustained from non-sustained attention periods within individuals (Seeburger et al., 2024). Their sensitivity to behavioral and cognitive states further supports their role in intrinsic functional dynamics, with studies showing that external stimuli, such as structured visual stimulation, can modulate QPP amplitude and phase (Xu, Smith, et al., 2023).

Complementing QPPs, cPCA provides an alternative framework for characterizing spatiotemporal dynamics by leveraging both amplitude and phase information to extract coherent, structured patterns from rs-fMRI data (Bolt et al., 2022, 2025). While QPPs iteratively refine a propagating template, cPCA decomposes signals in the complex domain, capturing phase-coherent fluctuations that shape network interactions. A direct comparison of these methods in the Human Connectome Project (HCP) dataset (Van Essen, Smith, et al., 2013) revealed that QPPs and cPCA extract highly similar network dynamics, suggesting that they capture overlapping intrinsic functional connectivity (Bolt et al., 2022). However, the use of HCP data, with its fast TR of 0.72s and approximately one hour of total scan time per participant across four sessions in a large corpus of participants, represents a high quality dataset and a pseudo-upper bound on what is feasible for clinical applications (as there are many exploratory and academic studies with shorter TRs). The extent to which these methods generalize to shorter scan durations and longer TRs, common in standard fMRI studies, remains an open question. Addressing these gaps is crucial for translating QPP and cPCA analyzes into more applied research and clinical contexts.

Despite their promise, it is not yet clear how sensitive the spatiotemporal patterns obtained from the QPP algorithm and cPCA are to methodological choices such as scan duration, repetition time (TR), and bandpass filtering. We therefore hypothesize that variation in these parameters has a substantial impact on the reproducibility of spatiotemporal patterns across scans and datasets. For example, longer scans may improve the stability of extracted patterns by capturing more recurrences of underlying network dynamics, but they may also introduce additional variability due to gradual changes in brain activity over time (Duda et al., 2023). In contrast, shorter scans are more susceptible to noise and individual-level variability (Birn et al., 2013; Laumann et al., 2015; Petersen et al., 2024), which can reduce the reliability of pattern detection. These methodological considerations motivate the present work, in which we systematically examine how parameter choices shape QPP- and cPCA-derived spatiotemporal templates and their test–retest reliability.

Similarly, TR influences how spatiotemporal patterns are sampled and represented. Shorter TRs offer finer temporal resolution, potentially improving sensitivity to fast dynamics, whereas longer TRs may reduce the accuracy of representing propagating patterns due to under-sampling. Notably, since the QPP algorithm uses TR to define the optimal window length for detection, TR inherently constrains the temporal resolution at which these patterns can be meaningfully extracted and interpreted. Finally, bandpass filtering plays a crucial role in shaping the frequency content of the data, with different filter settings highlighting distinct aspects of neural dynamics (Zalesky et al., 2014).

To investigate these methodological considerations, this study systematically evaluates how scan duration, TR, and bandpass filtering influence the stability and fidelity of spatiotemporal patterns extracted via the QPP algorithm and cPCA. Here, we use “fidelity” to refer to the consistency of the extracted spatial and temporal pattern structure under varying acquisition and preprocessing parameters, rather than agreement with an external ground truth, which is inherently unavailable in resting-state BOLD data. Specifically, we ask:

1. **Scan duration**: How does the amount of acquired data affect the stability and reproducibility of the extracted spatiotemporal patterns?
2. **TR selection**: How does sampling rate affect the detection of QPPs and cPCA-derived dynamics?
3. **Bandpass filtering**: How do different frequency bands influence the stability and temporal characteristics (e.g., timing and duration) of these patterns?

By directly comparing these methodological factors, we provide critical insights into the conditions under which QPP and cPCA yield reliable spatiotemporal templates. These findings offer practical guidance for optimizing spatiotemporal analysis in both basic neuroscience and clinical applications and contribute to establishing best practices for capturing dynamic functional connectivity with greater reproducibility across studies.

## 2 Methods

This study combines multi-dataset preprocessing, dual-level brain parcellation, and complementary spatiotemporal analysis algorithms to examine the reliability and fidelity of intrinsic BOLD signal dynamics.

As shown in Figure 1, we applied a standardized preprocessing pipeline to five independent resting-state fMRI datasets, mapped BOLD signals onto both region- and network-level parcellations, and analyzed spatiotemporal structure using two algorithms: Quasi-Periodic Patterns (QPPs) and complex Principal Component Analysis (cPCA). The following sections detail the data sources, preprocessing steps, algorithm implementations, and statistical validation procedures used in the study.

**Figure 1:**
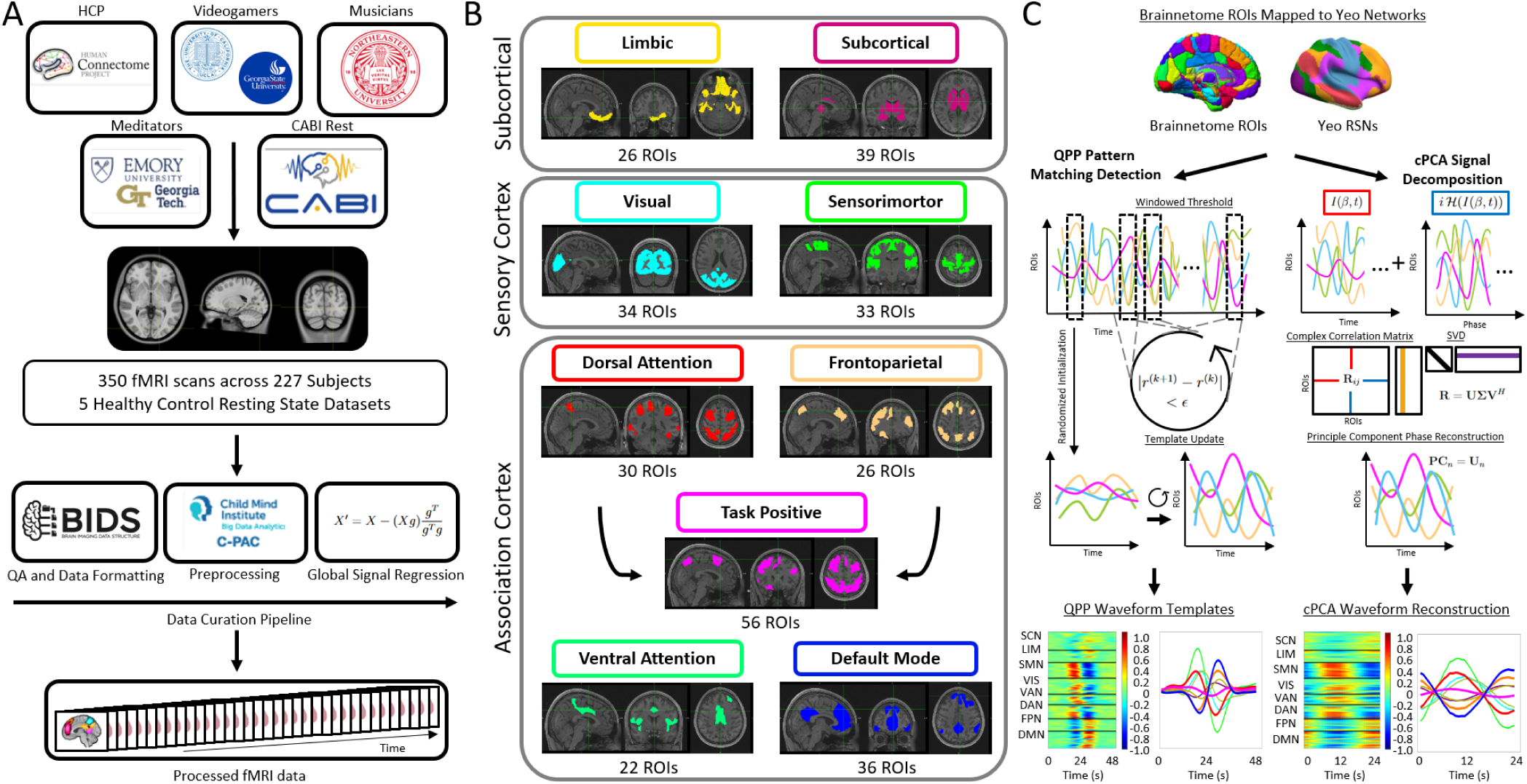
*Overview of the study design and methodology.* (A) Data preprocessing pipeline applied to all five datasets, including quality control for selecting viable scans, organization into the Brain Imaging Data Structure (BIDS), and preprocessing using the Configurable Pipeline for the Analysis of Connectomes (C-PAC) to ensure standardized neuroimaging preprocessing. All scans are global signal regressed. (B) Brain parcellation using the Brainnetome atlas, consisting of 246 regions of interest (ROIs) corresponding to the eight canonical Yeo networks: Default Mode (36 ROIs), Dorsal Attention (30 ROIs), Frontoparietal (26 ROIs), Ventral Attention (22 ROIs), Task Positive (56 ROIs), Visual (34 ROIs), somatomotor (33 ROIs), and Subcortical (39 ROIs). ROIs are grouped into three major cortical subdivisions: subcortical, sensory cortex, and association cortex. (C) Illustration of the spatiotemporal pattern detection algorithms used in this study: Quasi-Periodic Patterns (QPPs, left), and complex Principal Component Analysis (cPCA, right).

### 2.1 Data

rs-fMRI data were obtained from five independent datasets, totaling 227 participants across 350 scans. These datasets include a selection of the Human Connectome Project (HCP) (Van Essen, Smith, et al., 2013), a cohort of experienced videogamers acquired at Georgia State University (Videogamers) (Jordan & Dhamala, 2023), a dataset of trained musicians from the Northeastern Biomedical Imaging Center and the Olin Neuropsychiatry Research Center (Musicians) (Belden et al., 2020), a mindfulness meditation study from Emory University (Meditators) (Hasenkamp et al., 2012), and a cognitive task-based dataset from the Center for Advanced Brain Imaging (CABI) at Georgia Institute of Technology (CABI Rest) (Godwin et al., 2017). While some datasets included task-related fMRI scans, only the resting-state fMRI data were used in this study to systematically evaluate the algorithms across a diverse set of datasets and acquisition parameters. Participant demographics, including sex distribution, age (mean ± SD), number of scans per subject, total scan count, TR, and scan duration, are summarized in Table 2.

**Table 1:** List of Acronyms.

| Acronym | Definition |
| --- | --- |
| **Disorders / Cohorts** |  |
| AD | Alzheimer’s disease |
| ADHD | Attention-deficit/hyperactivity disorder |
| MCI | Mild cognitive impairment |
| NC | Normal controls |
| **Datasets / Study Labels** |  |
| ADNI | Alzheimer’s Disease Neuroimaging Initiative |
| CABI | Center for Advanced Brain Imaging |
| CABIRest | CABI resting-state dataset label (as used in tables/figures) |
| HCP | Human Connectome Project |
| Med | Meditators dataset label (as used in figures/appendix) |
| Music | Musicians dataset label (as used in figures/appendix) |
| VG | Videogamers dataset label (as used in figures/appendix) |
| **Imaging / Acquisition / Preprocessing** |  |
| 3T | 3 Tesla (scanner field strength) |
| BIDS | Brain Imaging Data Structure |
| BOLD | Blood-oxygen-level-dependent |
| EPI | Echo-planar imaging |
| FOV | Field of view |
| GSR | Global signal regression |
| IRB | Institutional review board |
| MNI | Montreal Neurological Institute (standard space) |
| MPRAGE | Magnetization-prepared rapid acquisition gradient echo |
| MRI | Magnetic resonance imaging |
| QC | Quality control |
| rs-fMRI | Resting-state functional magnetic resonance imaging |
| SNR | Signal-to-noise ratio |
| T1 | T1-weighted anatomical image |
| TE | Echo time |
| TR | Repetition time |
| **Parcellations** |  |
| dFC | Dynamic functional connectivity |
| FC | Functional connectivity |
| ICN | Intrinsic connectivity network |
| ROI | Region of interest |
| RSN | Resting-state network |
| **Canonical Network Abbreviations** |  |
| DAN | Dorsal Attention Network |
| DMN | Default Mode Network |
| FPN | Frontoparietal Network |
| LMN | Limbic Network |
| SCN | Subcortical Network |
| SMN | Somatomotor Network |
| TPN | Task-Positive Network (DAN & FPN combined) |
| VAN | Ventral Attention Network |
| VIS | Visual Network |
| **Methods / Algorithms** |  |
| cPCA | Complex principal component analysis |
| PC | Principal component |
| QPP | Quasi-periodic pattern |
| SVD | Singular value decomposition |
| SWC | Sliding-window correlation |
| **Frequency Bands / Filtering Labels** |  |
| Infraslow+ | Frequency band 0.01–0.15 Hz |
| Infraslow | Frequency band 0.01–0.10 Hz |
| Slow-5 | Frequency band 0.01–0.027 Hz |
| Slow-4 | Frequency band 0.027–0.073 Hz |
| **Pattern Alignment Metrics** |  |
| MJI | Modified Jaccard Index |
| MJI-C | Modified Jaccard Index with Cushion (temporal tolerance; paper-specific label) |
| PTPA | Precursor Time Point Analysis |
| TMX1 | Template maxima (instances of matched maxima; paper-specific label) |
| TMX2 | Template minima (instances of matched minima; paper-specific label) |
| **Statistics / Inference** |  |
| ANOVA | Analysis of variance |
| KW | Kruskal–Wallis test |
| MWU | Mann–Whitney U test |
| STD | Standard deviation |
| WSR | Wilcoxon signed-rank test |
| **Preprocessing Software** |  |
| C-PAC (CPAC) | Configurable Pipeline for the Analysis of Connectomes |

**Table 2:** Summary of fMRI datasets, including number of subjects, scan details, age statistics, repetition time (TR), and scan duration.

| Dataset | # Subjects (F/M) | # Scans/Subject | Total Scans | Age (Avg ± STD) | TR (secs) | Avg Scan Length<br>(Timepoints / Mins) | Total Group Length<br>(Timepoints / Hrs) |
| --- | --- | --- | --- | --- | --- | --- | --- |
| HCP | 20 (10/10) | 4 | 80 | 29.57 ± 3.8 | 0.72 | 1200 / 14.4 | 96,000 / 19.20 |
| Videogamers | 46 (16/30) | 4 | 168 | 20.36 ± 2.52 | 0.535 | 860 / 7.7 | 144,480 / 21.47 |
| Musicians | 48 (36/12) | 1 | 48 | 20.6 ± 2.4 | 0.475 | 937 / 7.4 | 44,979 / 5.93 |
| Meditators | 14 (11/3) | 2 | 14 | 46.4 ± 12.0 | 1.5 | 213 / 5.3 | 2,982 / 1.24 |
| CABI Rest | 99 (55/44) | 1-2 | 172 | 26 ± (7.0) | 2.25 | 146 / 5.5 | 25,112 / 15.70 |

Across all datasets, T1-weighted anatomical images were collected using a 3D magnetization-prepared rapid gradient-echo (MPRAGE) sequence on Siemens 3T MRI scanners, with voxel sizes ranging from 0.7 to 1 mm^3^. Functional scans were acquired using echo-planar imaging (EPI) sequences, with TRs ranging from 0.475 to 2.25 seconds and voxel resolutions between 2 and 4 mm^3^. The total scan duration varied between 5.3 and 14.4 minutes per run, with total group-level acquisition times ranging from 1.24 to 21.47 hours, depending on the dataset. A comprehensive overview of scan acquisition parameters, including field of view, flip angles, and slice counts, is provided in Supplementary Table S1.

All original scan acquisition adhered to best practices for ethical human data collection and informed consent in accordance with local institutional review boards. See original data acquisition study for details: HCP: Van Essen, Smith, et al., 2013; Videogamers: Jordan and Dhamala, 2023; Musicians: Belden et al., 2020; Meditators: Hasenkamp et al., 2012; and CABI Rest: Godwin et al., 2017.

### 2.2 Image Preprocessing

For processing, scans were formatted according to the Brain Imaging Data Structure (BIDS: https://bids.neuroimaging.io/, Poldrack et al., 2024). All preprocessing was conducted on a Linux (Ubuntu 22.04.3 LTS) system using the Configurable Pipeline for the Analysis of Connectomes (C-PAC: https://fcp-indi.github.io/ Craddock et al., 2013). Outputs were generated both with and without global signal regression (GSR). Only outputs with GSR were used for this study.

As part of the standard C-PAC pipeline, anatomical (T1-weighted) scans were registered to the 2mm Montreal Neurological Institute (MNI) 152 Atlas, ensuring alignment across subjects. rs-fMRI scans were subsequently registered to the same standard MNI space and extracted as time series to the Brainnetome 246 atlas, a parcellation scheme consisting of 246 regions of interest (ROIs) based on cytoarchitectonic and functional imaging (Fan et al., 2016).

In addition to the Brainnetome atlas, rs-fMRI signals were also mapped onto the eight functional networks defined by Yeo et al., 2011, which provide a large-scale functional organization of the cortex. Additionally, the Dorsal Attention Network (DAN) and Frontoparietal Network (FPN) were analyzed separately and also combined into a single Task-Positive Network (TPN), in alignment with previous psychology studies (Di & Biswal, 2014; Fox et al., 2005; Raichle et al., 2001; Seeburger et al., 2024; Watters et al., 2025). This approach follows the historical development of these network classifications: the TPN was originally defined as functionally opposing the Default Mode Network (DMN), with later refinements (e.g., Yeo’s parcellation) distinguishing DAN and FPN as distinct but overlapping subdivisions of the broader task-positive framework (Seeburger et al., 2024). This modification allows for an integrated analysis of attentional and executive control functions associated with these networks, facilitating a fidelity assessment of their combined influence. The networks were categorized into three major groups based on functional specialization: subcortical, sensory cortex, and association cortex. See Table 3 for further network-specific details.

**Table 3:** Functional network groups and their descriptions.

| Network Group | Abbreviation | Name | Description |
| --- | --- | --- | --- |
| Subcortical | SCN | Subcortical Network | Integration of subcortical structures related to affect, motivation, and autonomic control. |
|  | LBN | Limbic Network | Emotional and memory-related processing. |
| Sensory Cortex | SMN | Somatomotor Network | somatomotor integration and execution. |
|  | VIS | Visual Network | Primary and higher-order visual processing. |
| Association Cortex | VAN | Ventral Attention Network | Stimulus-driven attentional reorientation. |
|  | DAN | Dorsal Attention Network | Goal-directed attentional control. |
|  | FPN | Frontoparietal Network | Cognitive control and adaptive reasoning. |
|  | TPN | Task-Positive Network | Combination of DAN and FPN, reflecting goal-directed attentional control and cognitive flexibility. |
|  | DMN | Default Mode Network | Internally directed cognition and self-referential thought. |

By extracting time series from both the Brainnetome 246 atlas and Yeo’s eight functional networks, as well as incorporating the combined Task-Positive Network (TPN), we aim to examine network-level interactions while also resolving finer-grained regional dynamics. This dual-level parcellation enables a comprehensive analysis of functional connectivity, allowing comparisons between region-based and network-based approaches in resting-state fMRI.

### 2.3 Algorithms

**Quasi-Periodic Pattern Analysis (QPPs):** This algorithm identifies recurring spatiotemporal patterns in BOLD signal time series using a sliding window approach (Majeed et al., 2009). By iteratively updating a template based on high-correlation windows, the QPP algorithm refines its pattern characterization through a convergence process guided by correlation peaks. The process begins with selecting a random seed time point and computing a template over a defined window length. Sliding correlations with the BOLD fMRI time series are used to identify local maxima, which in turn update the template for subsequent iterations. Adjustments in thresholds and optional expansions of the window length further enhance pattern resolution (Xu, Yousefi, et al., 2023; Yousefi et al., 2018). For this study, a window length (WL) of 24 seconds was used, as this has been demonstrated to be an optimal duration for detecting QPPs (Maltbie et al., 2022). While the algorithm is capable of detecting multiple QPPs, the primary QPP following global signal regression has been the most extensively examined in prior studies (Daley et al., 2024; Watters, Fazili, Daley, Belden, LaGrow, et al., 2024; Watters et al., 2025). This dominant QPP, typically reflecting alternating BOLD amplitude modulations between the TPN and the DMN, serves as the main focus of this analysis. The final output consists of a refined template representing the QPP and its associated correlation peaks, providing a robust way to examine repetitive spontaneous whole-brain patterns. For the full algorithm, see Algorithm 1.

#### Algorithm 1

Quasi-Periodic Pattern Analysis (QPPs) for BOLD Signals

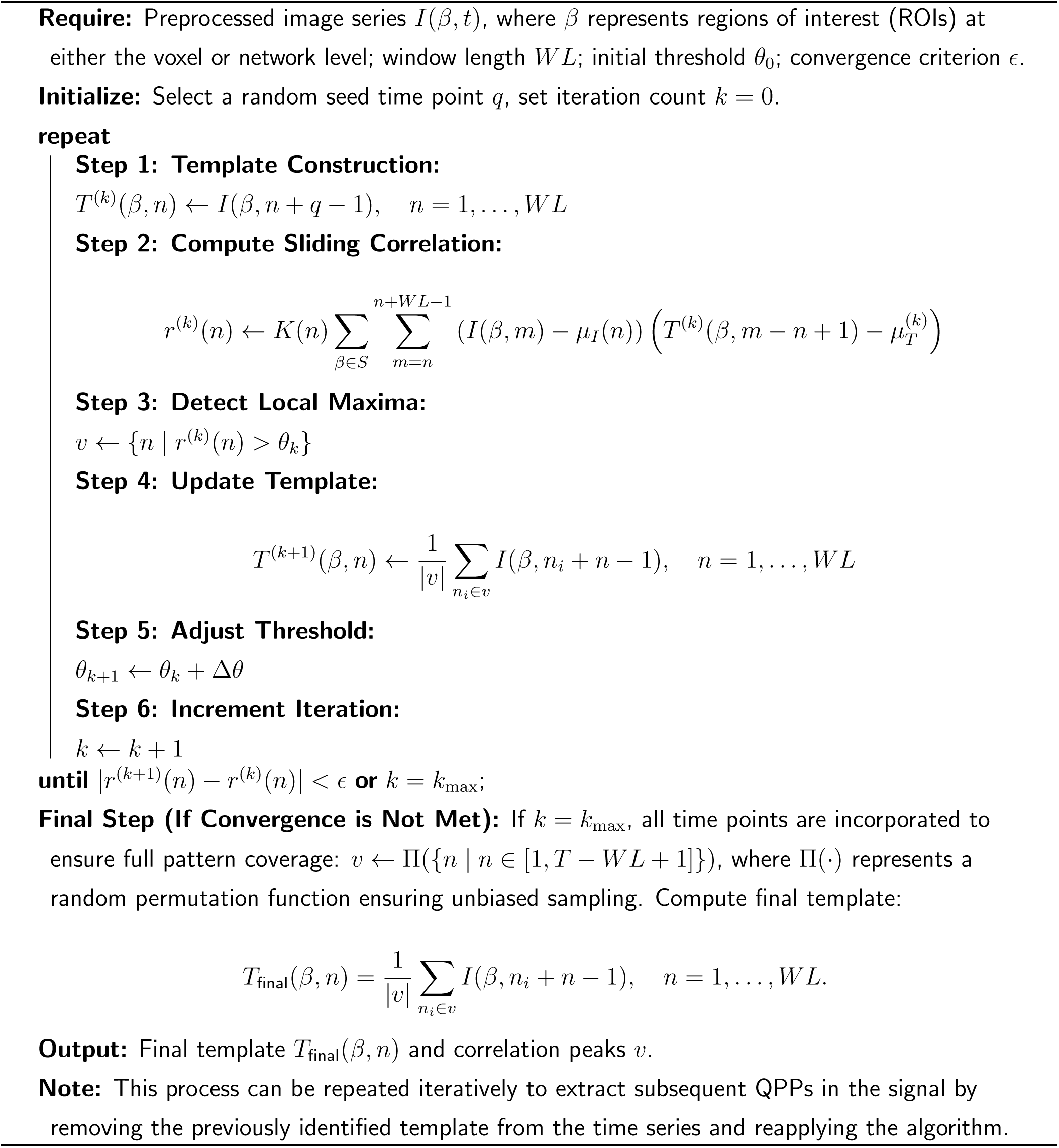

**Complex Principal Component Analysis (cPCA):** This algorithm extends standard principal component analysis to the complex domain, enabling the examination of both amplitude and phase information in BOLD signals. The cPCA approach begins with the application of the Hilbert transform to create complex-valued signal representations (Feeny, 2008; Horel, 1984). From these transformed signals, a complex correlation matrix is computed, capturing the strength of synchronization and phase relationships between brain regions. Singular value decomposition (SVD) is applied to extract the principal components, with only the top components retained for further analysis (Bolt et al., 2022). While multiple components can be extracted, we focus on the first principal component following GSR, as it has been the most widely examined in prior studies and is most comparable to the primary pattern extracted with the QPP algorithm. This component provides insight into the dominant phase-coherent spatiotemporal pattern, highlighting large-scale network interactions and their temporal shifts. Convergence is determined through changes in singular values, and the resulting phase delay maps offer critical insights into spatiotemporal phase variations within the data. This approach facilitates a deeper understanding of brain network dynamics, emphasizing phase relationships in neural activity. For the full algorithm, see Algorithm 2.

#### Algorithm 2

Complex Principal Component Analysis (cPCA) for BOLD Signals

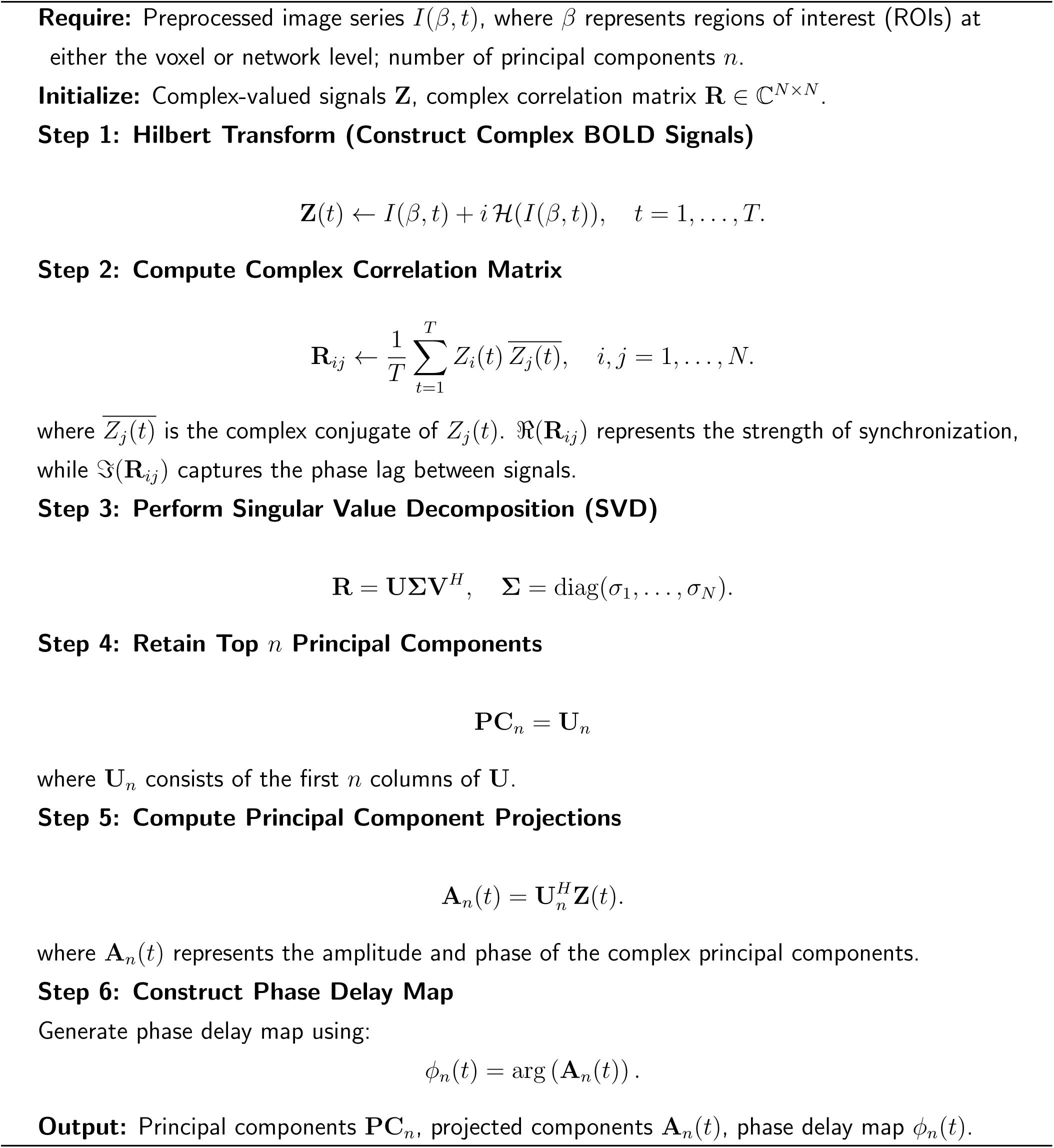

QPP and cPCA outputs were plotted as 246 ROIs assigned to Yeo’s canonical networks, which assigns cortical ROIs into 8 major association and sensory cortices, plus subcortical regions (See Supplementary Table S2 in the appendix for a key used to group Brainnetome ROIs into Yeo’s networks).

The following networks were used: subcortical network (SCN), limbic network (LIM), somatomotor network (SMN), visual network (VIS), ventral attention network (VAN), dorsal attention network (DAN), frontoparietal network (FPN), task positive network (TPN), and default mode network (DMN). As indicated in the following sections, the primary factor driving the anticorrelation between the DMN across brain states appears to be DMN/DAN anticorrelation, as shown in previous studies (Seeburger et al., 2024; Watters, Fazili, Daley, Belden, LaGrow, et al., 2024). QPP dynamics for the major attentional and sensory networks have been extensively examined at various levels of analysis (Watters, Fazili, Daley, Belden, LaGrow, et al., 2024; Watters et al., 2025); however, limbic and subcortical networks are often excluded. Subcortical systems, in particular, are known to participate in whole-brain spatiotemporal patterns and to anchor large-scale propagation motifs (Yousefi & Keilholz, 2021). This is one of the first studies to report the spatiotemporal patterns of the subcortical networks across a diverse set of data.

### 2.4 Statistical and Validation Procedures

#### 2.4.1 Assessing Individual and Group Analysis

To compare spatiotemporal patterns extracted from typical and HCP-quality rs-fMRI data, we analyzed two large-scale resting-state fMRI datasets: HCP and CABI Rest (see Table 2 and Supplementary Table S1). These datasets were selected because they provide both multiple scanning sessions and varying scan durations, enabling a systematic investigation of how data quantity and sampling characteristics impact the performance of QPP and cPCA algorithms.

The HCP dataset includes four resting-state fMRI sessions per subject, each with a scan duration of approximately 15 minutes and a fast temporal resolution (TR = 0.72 seconds), facilitating fine-grained temporal analyses. In contrast, the CABI Rest dataset includes up to two sessions per subject, each with a scan duration of approximately 7 minutes and a lower temporal resolution (TR = 2.25 seconds). This allows us to assess how both reduced scan duration and coarser temporal sampling affect the reliability of the extracted spatiotemporal patterns.

By comparing these datasets, we evaluate the feasibility of individual-subject analyses under more typical clinical-quality acquisitions (e.g., ADNI, CABI) and contrast this with group-level analyses applied to high-quality, high-quantity data (HCP), where longer, low-TR scans and multiple sessions allow averaging across subjects to mitigate noise and stabilize the extracted patterns. The dataset characteristics and preprocessing details are summarized in Appendix Table A1.

For individual-subject analysis, we set one hour of total scan time per subject as the practical upper bound, reflecting real-world constraints in fMRI research. Longer sessions are often impractical due to cost constraints (Asnafi et al., 2020), limited scanner availability (Nakagawa et al., 2013), and participant fatigue (Lieberman & Cunningham, 2009). Furthermore, studies requiring repeated scans per subject face logistical barriers, particularly in clinical and developmental populations where extended scanning sessions may not be feasible. Given that HCP provides a maximum of four sessions per subject, and CABI Rest provides a maximum of two, we aimed to assess whether meaningful spatiotemporal patterns could be extracted from a single scan or an accumulated scan time of up to one hour per subject.

For group-level analysis, we considered scan durations exceeding one hour, reflecting common aggregation approaches in large-scale neuroimaging studies. Public datasets frequently compile tens to hundreds of hours of fMRI data to enhance statistical power and generalizability (e.g., HCP (Van Essen, Ugurbil, et al., 2013), OpenNeuro (Castrillon et al., 2022), UK Biobank (Miller et al., 2016), BigBrain (Amunts et al., 2013), to name a few). To ensure comparability across methods, we set the upper bound for analysis at 14.4 hours, aligning with the total available scan time in the CABI Rest dataset. This allowed us to systematically examine how algorithmic performance scales with increasing data availability.

To assess the fidelity of extracted spatiotemporal patterns, we evaluated the correlation between the DMN and other canonical Yeo networks using Pearson correlation. The DMN is a core intrinsic network involved in memory, self-referential thought, and consciousness, with activity predominating during wakeful rest and negatively correlated with externally directed attention (Andrews-Hanna et al., 2010; Buckner et al., 2008; Fox et al., 2005; Greicius et al., 2003; Raichle et al., 2001). Given its fundamental role in intrinsic brain activity, its correlation with other large-scale networks serves as a benchmark for assessing the stability and biological relevance of extracted patterns across different scan durations and analysis levels. By comparing individual- and group-level analyses, we aimed to determine whether these methods reliably capture reproducible signatures of brain activity and how their application should vary for individual inference versus population-level modeling.

To statistically assess differences in functional connectivity patterns, we employed the Kruskal-Wallis test, a non-parametric method suitable for comparing distributions without assuming normality. This approach is particularly appropriate for functional connectivity data, which often exhibits non-Gaussian distributions and unequal variances across groups, making it more robust than one-way ANOVA, which assumes normality (Couch et al., 2019). A significance threshold of *α* = 0.05 was used.

For pairwise comparisons, we applied the Wilcoxon signed-rank test for within-dataset analyses and the Mann-Whitney U test for between-group comparisons. The Wilcoxon signed-rank test is a non-parametric alternative to the paired t-test, suitable for assessing differences in functional connectivity. The Mann-Whitney U test, on the other hand, serves as a robust alternative to the independent t-test, allowing us to compare distributions between separate groups without assuming normality.

When the Kruskal-Wallis test indicated a significant difference across groups, Dunn’s test was applied as a post-hoc analysis to identify which specific group comparisons were significant (Dinno, 2015). To rigorously control for Type I errors, Dunn’s test included a Bonferroni correction for multiple comparisons. This statistical framework ensures that observed differences in functional connectivity are not artifacts of non-normal data distributions, thereby enhancing the validity and interpretability of our findings.

#### 2.4.2 Effect of Repetition Time (TR) on Analysis

To evaluate how different TRs affect QPPs and cPCA, we systematically resampled the TR across datasets. The simplest method of synthetically increasing TR for faster-sampled datasets was by selecting every second, third, or subsequent time point, effectively increasing the TR by integer multiples of the original value. For instance, in the HCP dataset, the native TR of 0.72s was resampled to 1.44s, 2.16s, 2.88s, and 3.60s. Although there could be benefits in exploring even longer TRs, we focused on resampling high-temporal-resolution datasets rather than interpolating lower-resolution data, as the latter would introduce additional assumptions about signal continuity and could obscure the native temporal structure of QPPs and cPCA components.

To ensure independent comparisons across TR values, each dataset was divided into non-overlapping groups with independent subjects and scans, maintaining approximately equal total scan time within each group. The available data included a sufficient number of subjects and sessions to construct multiple groups with distinct subjects, while still retaining multiple scans per subject within each group. This structure enabled within-group averaging to improve reliability, while preserving independence across groups for comparison. This design allows for a controlled assessment of how TR influences QPP detection and cPCA-derived components across both fast- and slow-sampled datasets.

The delineations for each dataset, including the number of groups, subjects per group, scans per group, corresponding TR values, and total scan time per group, are summarized in Table 4.

**Table 4:** Repetition time (TR) delineations across datasets. TR was systematically increased by resampling, allowing for independent comparisons across different sampling rates. Scan time per group represents the total duration of data used for each delineation.

| Dataset | Groups | Subjects /Group | Scans /Group | Scan Time /Group (hrs) | TR (s) | Resampled TRs (s) |
| --- | --- | --- | --- | --- | --- | --- |
| HCP | 5 | 4 | 16 | 3.84 | 0.72 | 1.44, 2.16, 2.88, 3.60 |
| Videogamers | 6 | 7 | 28 | 3.58 | 0.535 | 1.070, 1.605, 2.140, 2.675, 3.210 |
| Musicians | 7 | 6 | 6 | 0.74 | 0.475 | 0.950, 1.425, 1.900, 2.375, 2.850, 3.325 |
| Meditators | 2 | 7 | 7 | 0.62 | 1.5 | 3.0 |
| CABI Rest | 1 | 20 | 40 | 3.65 | 2.25 | n/a |

#### 2.4.3 Frequency’s Significance

rs-fMRI studies often employ bandpass filtering to isolate relevant frequency components of the BOLD signal. rs- fMRI encompasses both neural and physiological components of the BOLD signal, including infraslow neural oscillations and physiological artifacts such as cardiac and respiratory fluctuations (Birn et al., 2006; Biswal et al., 1995; Bolt et al., 2024; Cordes et al., 2001). Within neurologically and physiologically relevant frequencies, the infraslow frequency band (0.01-0.1 Hz) is the most commonly used for rs-fMRI analysis and defaulted on widely used preprocessing pipelines such as the Configurable Pipeline for the Analysis of Connectomes (C-PAC) Craddock et al., 2013. This passband is associated with spontaneous neural activity and functional connectivity and has been further subdivided based on neurophysiological studies into more distinct frequency bands (Gohel & Biswal, 2015; Lv et al., 2018).

The Slow-5 band (0.01-0.027 Hz) has been linked to fluctuations in arterial partial pressure of CO_2_ and O_2_, which in turn modulate the BOLD signal (Lv et al., 2018). This band may also reflect metabolic and vascular processes (Gohel & Biswal, 2015). The Slow-4 band (0.027-0.073 Hz) is considered to be more specific to neural activity and functional connectivity than Slow-5 (Gohel & Biswal, 2015). It has also been suggested to relate to the propagation of infraslow intrinsic BOLD signals within the brain (Betta et al., 2020).

Higher frequency bands, such as Slow-3 (0.073–0.198 Hz), have been less explored in rs-fMRI due to the inherent low sampling frequency of fMRI acquisitions. Nonetheless, studies suggest that this band has neurophysiological origins and functional significance, potentially reflecting synchronization of neuronal populations and modulation of cortical excitability (Gohel & Biswal, 2015). Similarly, the Slow-2 band (0.198–0.25 Hz) remains underexplored but has been proposed to relate to neuronal oscillations and cognitive processes, while also being strongly influenced by physiological noise, including cardiac and respiratory cycles (Gohel & Biswal, 2015) (see Figure 7A for an illustrative example). In the present study, we therefore restrict our analyses to infraslow, Slow-5, and Slow-4 bands, where physiological contamination is comparatively reduced and the dominant contributions are more clearly linked to canonical rs-fMRI networks.

In this study, we focus on four specific frequency bands to systematically evaluate their contributions to resting-state BOLD fluctuations: Slow-5 (0.01-0.027 Hz) and Slow-4 (0.027-0.073 Hz), which are implicated in infraslow oscillations, as well as two broader infraslow bands (0.01-0.10 Hz and 0.01-0.15 Hz). The latter two bands are included to test the extent to which broader spectral components influence observed patterns. The infraslow passband (0.01-0.15 Hz), denoted as ”Infraslow+” throughout the rest of the study, provides an opportunity to assess whether higher frequencies within the infraslow range contribute dominantly to signal variability. Both are widely used by previous studies (Iraji et al., 2022; Jensen et al., 2024; Lewis et al., 2023; Maltbie et al., 2022; Xu, Smith, et al., 2023; Yousefi & Keilholz, 2021). By comparing these four bandpass conditions, we aim to better characterize how different frequency components shape spatiotemporal analysis and functional connectivity in rs-fMRI.

#### 2.4.4 Comparing Instances of Spatiotemporal Patterns in the Datasets

To quantify the similarity of detected patterns, we employ a Modified Jaccard Index with Cushion, which extends the traditional Jaccard Index by incorporating a temporal tolerance, allowing for small deviations in timepoint alignment. This approach has been used in prior work to evaluate the robustness of neural event detection in time series data (Zhou & Wieser, 2019). Additionally, to assess temporal shifts between matched extrema across frequency bands, we implement a Precursor Time Point Analysis, which calculates the mean time differences between corresponding maxima and minima in different frequency bands within a predefined tolerance window. For further details on the derivations, please see the appendix. By integrating these two approaches, we aim to determine the degree of temporal alignment of the spatiotemporal patterns across different frequency band representations of the BOLD signal and evaluate whether cyclic activity exhibits coherence across conditions.

## 3 Results

Our first objective was to determine whether QPP and cPCA capture comparable spatiotemporal patterns across diverse datasets, and to evaluate the consistency of large-scale network dynamics revealed by each method. This initial analysis focused on comparing results across five distinct resting-state fMRI datasets (HCP, Videogamers, Musicians, Meditators, and CABI Rest), to assess whether both approaches recover similar representations of brain organization despite differences in data characteristics and processing pipelines.

Figure 2 and Figure 3 compare QPP- and cPCA-derived spatiotemporal patterns, showing that both methods capture similar large-scale network dynamics. Figures 2 and 3 further illustrate the differences in correlation structure between QPPs and cPCA across datasets, at both the Brainnetome ROI level and the Yeo network level. In Figure 2, panel A shows the extracted templates from both methods, demonstrating broadly consistent spatiotemporal patterns across algorithms, while still allowing for dataset-specific differences. Note that the x-axis scale differs between QPP and cPCA waveforms; the QPP algorithm uses a window twice as long to account for the extended temporal structure (“long-tailedness”) of its matched-filter approach. However, the central waveform extracted by both methods spans the same core duration, allowing for valid comparisons of temporal profiles. Panel B presents correlation matrices for the 246 Brainnetome ROIs, allowing direct comparison between QPP- and cPCA-derived connectivity. The difference matrices in panel C highlight methodological variations, with warm and cool colors indicating stronger correlations in cPCA and QPPs, respectively. Despite these differences, the overall patterns are highly similar, suggesting that both approaches robustly capture shared features of intrinsic functional organization.

**Figure 2:**
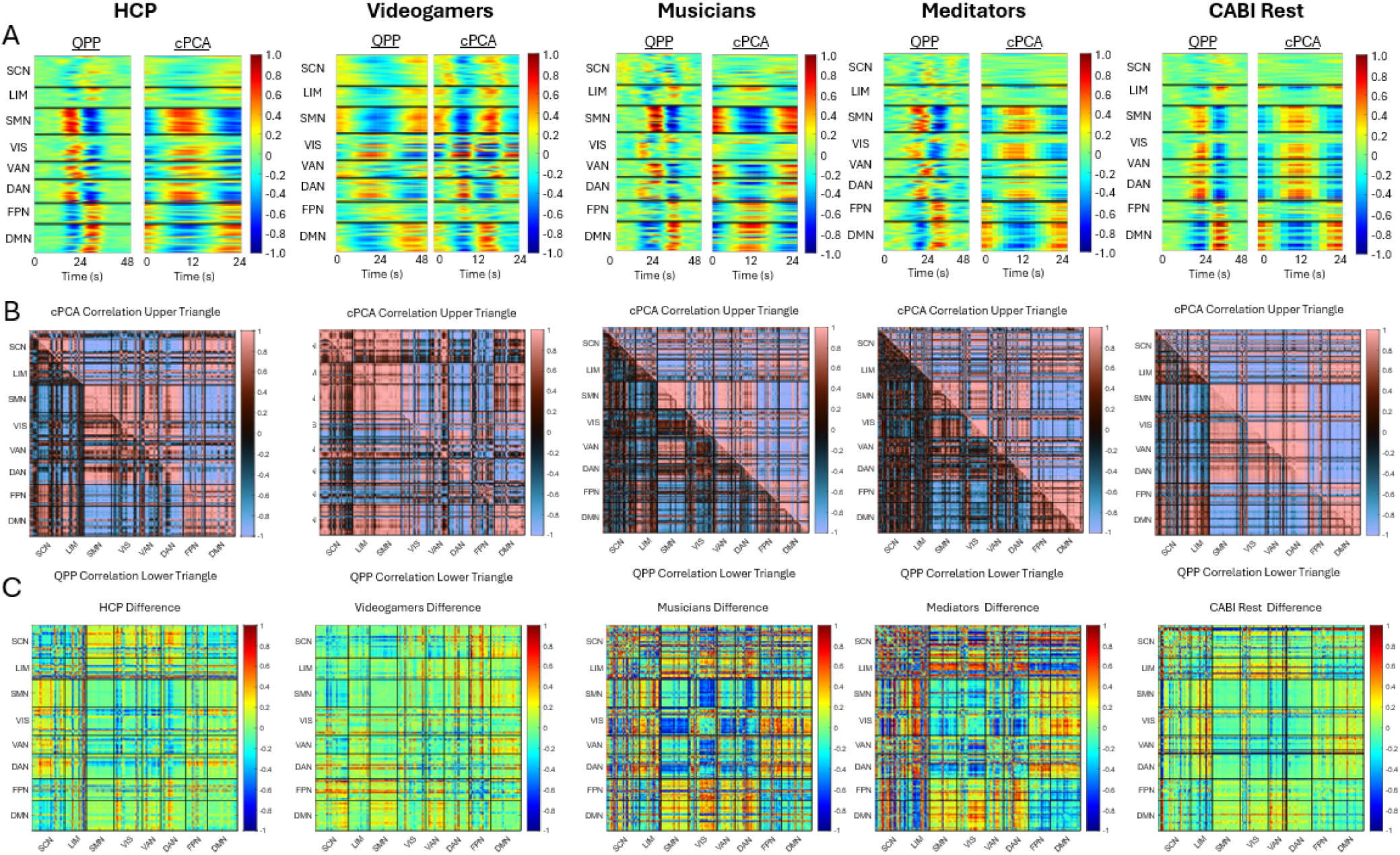
*Difference of Brainnetome ROI Correlation Between QPPs and cPCA Across Datasets*. (A) QPP and cPCA template waveforms extracted from each of the five datasets (HCP, Videogamers, Musicians, Meditators, and CABI Rest), plotted across the eight canonical Yeo networks: subcortical network (SCN), limbic network (LMN), somatomotor network (SMN), visual network (VIS), ventral attention network (VAN), dorsal attention network (DAN), frontoparietal network (FPN), and default mode network (DMN). Note that the x-axis scale differs between QPP and cPCA waveforms; the QPP algorithm uses a window twice as long to account for the extended temporal structure (”long-tailedness”) of its matched-filter approach. However, the central waveform extracted by both methods spans the same core duration, allowing for valid comparisons of temporal profiles. (B) Correlation matrices depicting the functional connectivity of each of the 246 Brainnetome ROIs. The lower triangular section of each matrix represents QPP-derived correlations, while the upper triangular section corresponds to cPCA-derived correlations, allowing for direct comparison across datasets. (C) Difference matrices showing the subtraction of QPP-derived correlations from cPCA-derived correlations for each dataset. Warmer colors indicate higher correlations in cPCA compared to QPPs, while cooler colors indicate higher correlations in QPPs relative to cPCA. These difference maps highlight methodological distinctions in network-level interactions captured by each approach.

**Figure 3:**
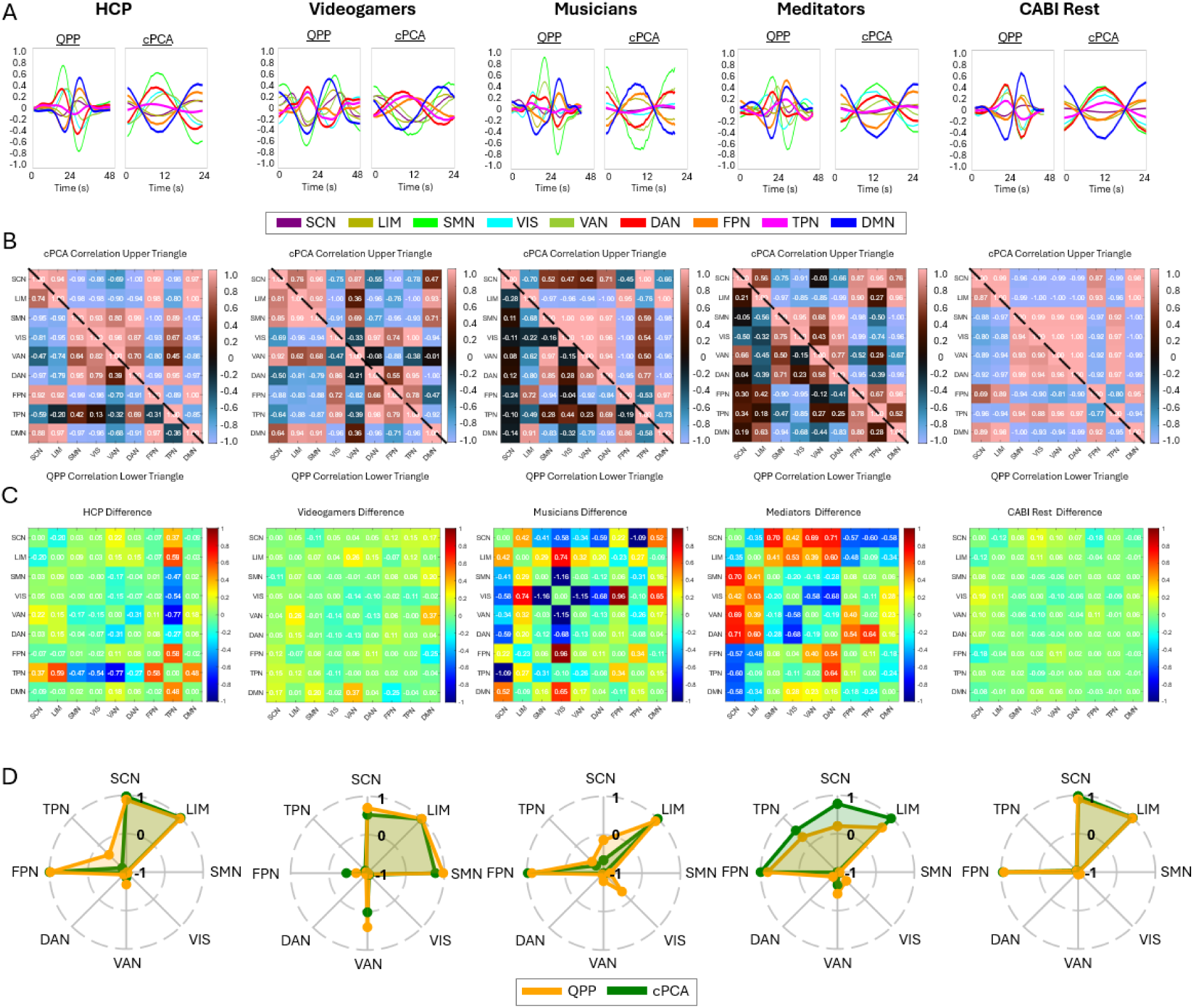
*Difference of Yeo Network Correlation Between QPPs and cPCA Across Datasets*. (A) QPP and cPCA template waveforms extracted from each of the five datasets (HCP, Videogamers, Musicians, Meditators, and CABI Rest), plotted across the eight canonical Yeo networks: subcortical network (SCN), limbic network (LMN), somatomotor network (SMN), visual network (VIS), ventral attention network (VAN), dorsal attention network (DAN), frontoparietal network (FPN), and default mode network (DMN). These waveforms illustrate the spatiotemporal progression of each method’s extracted patterns. Note that while QPP and cPCA often capture similar spatial patterns, their temporal alignment may differ in phase due to differences in how each method models propagation dynamics. (B) Correlation matrices depicting the functional connectivity between Yeo network-level time series. The lower triangular section represents QPP-derived correlations, while the upper triangular section corresponds to cPCA-derived correlations, facilitating direct comparison across datasets. (C) Difference matrices showing the subtraction of QPP-derived correlations from cPCA-derived correlations for each dataset. Warmer colors indicate higher correlations in cPCA compared to QPPs, while cooler colors indicate higher correlations in QPPs relative to cPCA. Notably, the Musicians and Meditators datasets show larger differences between methods, which may be partially attributed to the shorter total scan time available for each group (less than one hour), in contrast to the 3.5–3.8 hours available for the other datasets. This reduced scan time resulted from the need to subdivide these datasets into more granular groups to accommodate resampled TR conditions. These difference maps highlight methodological distinctions in network-level interactions captured by each approach. (D) Polar plots of the correlation between the Default Mode Network (DMN) and all other Yeo networks, with QPP (yellow) and cPCA (green) plotted for comparison. Correlation values range from −1 (center) to 1 (outermost edge).

Similarly, Figure 3 follows the same structure at the Yeo network level, revealing that while both methods capture large-scale network interactions, their relative strengths in inter-network connectivity differ. In

Figure 3, panel D shows polar plots of DMN correlations, revealing overall consistency between methods but with subtle differences in correlation strength and direction. Across all datasets, a consistent pattern emerges, showing strong agreement in broad network structures, with divergence limited primarily to finer-scale inter-network coupling.

### 3.1 Objective 1: Scan Length for Spatiotemporal Pattern Detection

We systematically assess how scan duration impacts the stability of QPP- and cPCA-derived connectivity patterns. This analysis provides insights into the feasibility of these methods for individual- and group-level studies, highlighting their relative robustness in capturing dynamic network interactions over time.

Figure 4 examines the relationship between scan length and DMN correlation for both QPP and cPCA-derived templates, providing insights into their stability and variability over time. Panels (A) and (B) display results separately for the HCP and CABI Rest datasets, which differ in TR: HCP was acquired with a TR of 0.72 seconds, offering fast temporal precision, while CABI Rest was acquired with a longer TR of 2.25 seconds, resulting in coarser temporal sampling. The top row illustrates how DMN correlation evolves with increasing scan duration, while the bottom row quantifies the variability in network correlations across independent subject groups. For a demonstration of linear x-axis across group scan length from 0–15 hours, please see Appendix Figure A1. For information on specific scan time length groups and number of instances, please see Appendix Table A1. A key finding is that differences between cPCA and QPPs vary across both scan length and network type. Notably, subcortical, sensory, and association cortices exhibit distinct patterns of variability in relation to DMN correlation.

**Figure 4:**
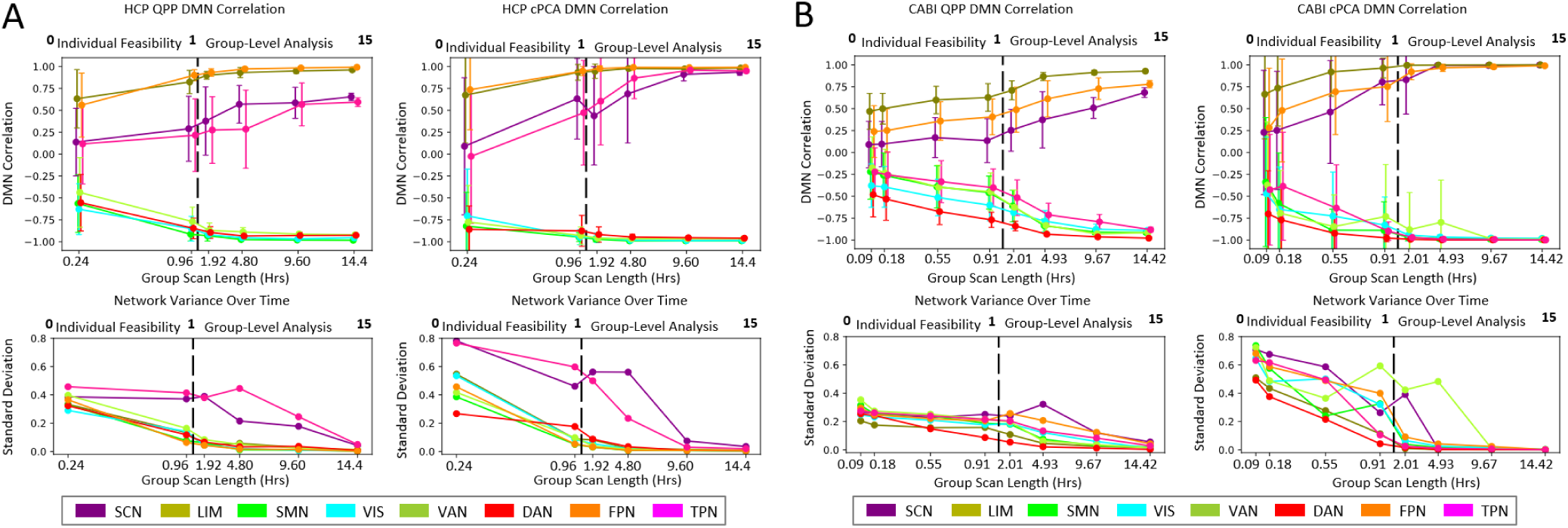
*DMN Correlation to Other Networks and Variability Over Scan Length for QPP and cPCA Analyses*. (A, B) DMN Correlation and Network Variability Over Time. The relationship between the DMN correlation and scan length is assessed for both QPPs and cPCA across two datasets: (A) HCP and (B) CABI Rest. Independent subject groups are used to evaluate variability, with the top row displaying DMN correlation over time and the bottom row showing the standard deviation of network variance over time. At 1 hour delineation, the black dashed line marks the methodological transition from individual feasibility to group-level analysis.

In the subcortical areas, variability in SCN-to-DMN correlation remains consistently high across scan lengths, indicating dynamic and heterogeneous contributions to DMN connectivity. This variability may also reflect lower signal-to-noise ratio (SNR) in subcortical regions, which are often less optimally captured by standard head coil configurations. Despite increased group-level averaging, subcortical correlation does not stabilize, suggesting a persistent influence of subject-level variability or acquisition-related limitations.

In contrast, the limbic network (LIM) maintains a relatively stable correlation with the DMN, exhibiting consistency across both algorithms and scan durations. Although this may seem unexpected if SNR were the primary driver of variability, it suggests that LIM-DMN coupling may be more reliable to both methodological and physiological sources of noise.

In the sensory cortex, the VIS and SMN remained consistently anticorrelated with the DMN across both individual feasibility and group-level analyses. This pattern was observed across both QPP and cPCA methods. However, variability within these networks was lower for QPPs compared to cPCA, particularly for higher TR datasets such as CABI Rest. This suggests that QPPs may provide a more stable characterization of sensory networks, particularly in shorter scans with shorter TR.

For the association cortex, results indicate consistent trends across scan lengths and algorithms. The task-positive network (TPN) remains highly variable, showing strong anticorrelation with the DMN, whereas the dorsal attention network (DAN) exhibits greater stability across both methods. These findings highlight that QPP and cPCA may differ in how they capture association network dynamics: QPP-derived measures tend to show lower variability across scan lengths, which may facilitate interpretation at the individual level by yielding more stable templates. In contrast, the higher variability observed in cPCA-derived measures may reflect greater sensitivity to inter-subject differences, capturing individual heterogeneity in network expression.

Furthermore, the standard deviation of DMN correlation derived from cPCA remains consistently higher across both datasets, suggesting that cPCA is more sensitive to inter-subject variability and may be less stable over shorter scan durations. In contrast, QPP-derived correlations exhibit lower variability, particularly in short scan lengths, implying a more stable and reproducible characterization of functional connectivity.

To better understand the practical implications of these methods, Appendix Figure A2 and Appendix Table A2 compare the computational complexity and processing times of QPP and cPCA across different scan lengths. QPP scales linearly with scan length, largely influenced by its sensitivity to window length (*WL*). In contrast, cPCA follows a quadratic complexity pattern with respect to the number of voxels (*N* ), making it theoretically more computationally intensive at larger scales. However, despite this higher theoretical complexity, cPCA remains computationally feasible for standard fMRI datasets, as illustrated in Appendix Figure A2.

For individual scans—such as CABI (146 timepoints) and HCP (1200 timepoints)—both QPP and cPCA are efficient and manageable. Even at the upper bound of individual feasibility (292 timepoints for CABI, 4800 for HCP), both methods remain practical for neuroimaging workflows. However, at the group level, where datasets can reach 25,112 timepoints for CABI or 96,000 for HCP, cPCA becomes significantly more demanding. Yet, optimized singular value decomposition (SVD) implementations help maintain its feasibility in large-scale analyses.

These findings suggest that QPPs may be more suitable for individual-level analyses, as its reduced variability allows for more reliable subject-specific characterization, whereas cPCA’s persistent variability may indicate greater suitability for group-level inferences where inter-subject differences are averaged out over longer durations.

### 3.2 Objective 2: TR’s Role in Spatiotemporal Fidelity

Repetition time (TR), the time between successive acquisitions of whole, brain volumes—sets the temporal resolution of the fMRI time series and therefore constrains the timescales of spatiotemporal dynamics that can be recovered. Longer TRs provide fewer samples per cycle of a transient pattern, which can blur waveform structure and reduce sensitivity to QPP and phase-coherent cPCA modes.

To systematically evaluate these effects across diverse datasets, Figures 5, A3, A4, and A5 (see Appendix) present resampled analyses across multiple TRs. In datasets with long total scan duration (approximately 3.5 hours of resting-state data per condition for HCP, Videogamers, and CABI Rest), both QPP and cPCA consistently preserved the dominant spatiotemporal patterns even when the data were resampled to longer TRs. This suggests that, given sufficient total scan length, the dominant patterns are relatively robust to moderate changes in temporal sampling at the level of aggregated (group-averaged) templates. At the same time, Figure 5E shows that specific network relationships do change with resampling: DMN–TPN coupling differs at the shortest TR, DMN–SCN correlations diminish at the longest TRs, and QPP-and cPCA-derived DMN connectivity estimates diverge more as TR increases. Thus, while the core spatiotemporal templates remain similar, finer-grained inter-network coupling measures are more sensitive to temporal resampling. Figure 5, for example, shows that despite resampling the HCP dataset from 0.72 s to 3.60 s, the extracted QPP and cPCA templates retain their core structural features across the eight Yeo networks.

**Figure 5:**
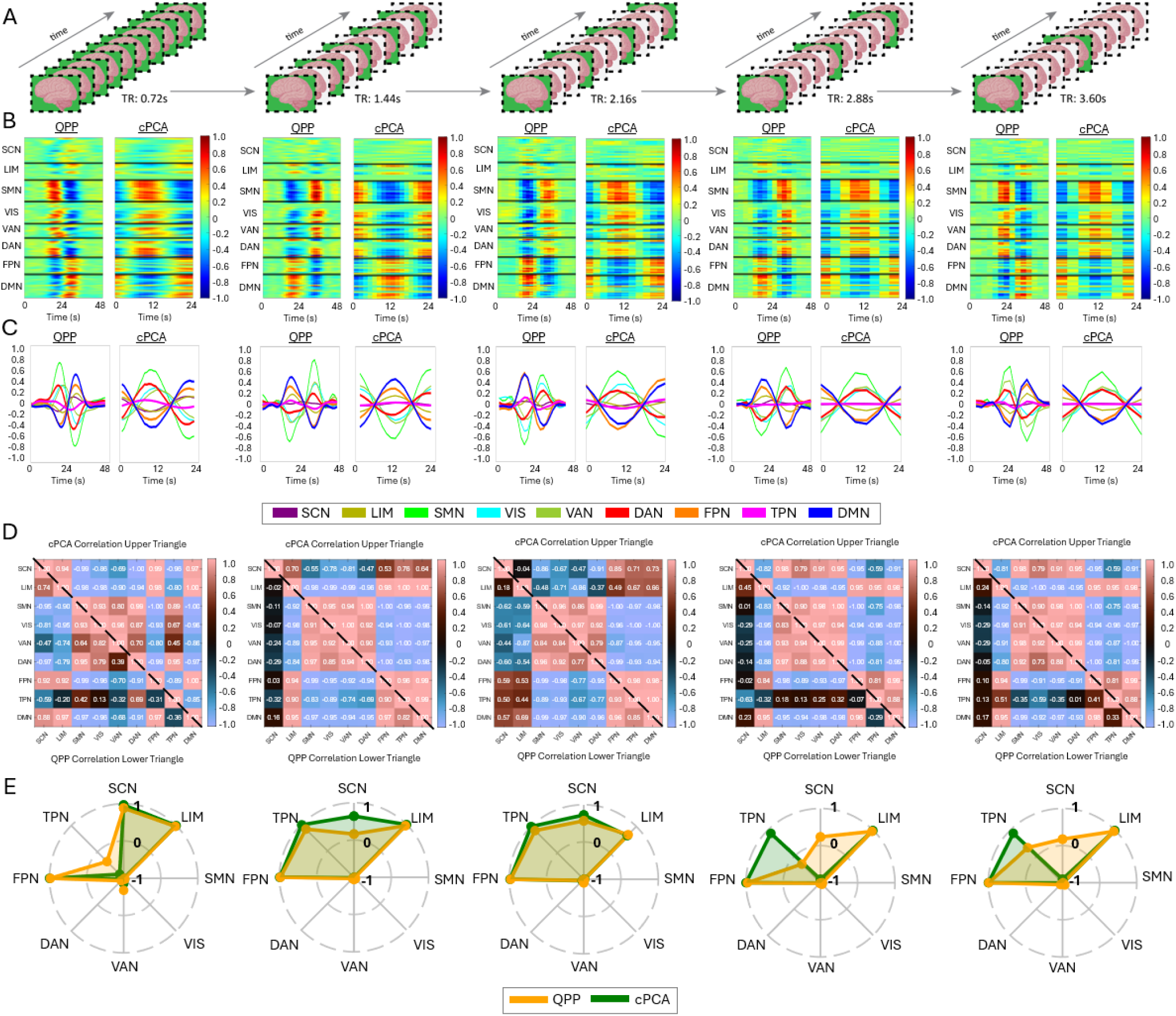
*Impact of Temporal Resampling on QPP and cPCA Detection in HCP Data.* (A) Illustration of five groups of resampled HCP fMRI data, demonstrating the effect of increasing TR. TR values range from 0.72s to 3.60s, allowing for systematic assessment of temporal resolution effects. (B-C) QPP and cPCA template waveforms extracted from each resampled dataset, plotted across the (B) 246 Brainnetome ROIs and (C) eight canonical Yeo networks (SCN, LMN, SMN, VIS, VAN, DAN, FPN, TPN, DMN). The overall waveform structure remains consistent across resampling conditions, though subtle variations emerge with increased TR. (D) Correlation matrices comparing QPP-derived and cPCA-derived templates, with the lower triangular matrix representing QPP correlations and the upper triangular matrix representing cPCA correlations. (E) Polar plots depicting the correlation between the default mode network (DMN) and all other Yeo networks. QPP-derived (yellow) and cPCA-derived (green) correlations are displayed for comparison, showing that the network dynamics between the DMN and the other networks remains preserved even at a resampled TR of 3.60s. These findings highlight the robustness of QPPs in capturing large-scale spatiotemporal dynamics despite reduced temporal resolution, while also demonstrating the sensitivity of cPCA-based templates to varying sampling rates.

It is important to note, however, that resampling inherently does not reconstruct the underlying neural signal as it would have been sampled at a higher TR; rather, it effectively subsamples the original time series, which may mask finer temporal details or introduce interpolation effects. Nevertheless, the observed stability in extracted patterns across resampled datasets—also evident in Appendix Figures A3 and A5 for the Videogamers and CABI Rest datasets—suggests that key network dynamics, particularly DMN connectivity patterns, remain robust to changes in TR.

In contrast, datasets with shorter scan durations—Musicians and Meditators, both with total group scan times under one hour—provide insight into the feasibility of these approaches at the individual-subject level. Figures A4 and A5 demonstrate that even at degraded TRs (3.325 s for Musicians and 3.00 s for Meditators), both QPP and cPCA continue to capture meaningful large-scale functional dynamics, with the dominant spatiotemporal trajectory and its primary network relationships remaining recognizable across resampling conditions. At the same time, the strength and balance of specific inter-network correlations do change with TR, indicating that finer-grained coupling estimates are more sensitive to temporal resolution than the overall pattern structure. This suggests that even within the constraints of limited scan duration and coarser TR, these methods can still reliably detect dominant network interactions at the individual level, while more subtle network relationships should be interpreted with greater caution.

When examining the polar plots across datasets (Figure 3 for cross-dataset comparison and Figure 5, Appendix Figure A3, Appendix Figure A4, Appendix Figure A5A, Appendix Figure A5B for individual datasets across resampled TRs), we observe both consistencies and variations. While QPP and cPCA exhibit substantial overlap in their extracted network templates, variability is evident in the SCN, FPN, and TPN across datasets and methods. These differences likely reflect the varying sensitivity of the methods to temporal resolution changes and their ability to capture transient network dynamics. However, when aggregated in Figure 6, the overall consistency in DMN interactions across datasets further supports the robustness of both methods. These findings highlight that, while temporal resampling influences fine-grained network dynamics, QPP and cPCA reliably capture dominant connectivity patterns across a range of temporal resolutions.

**Figure 6:**
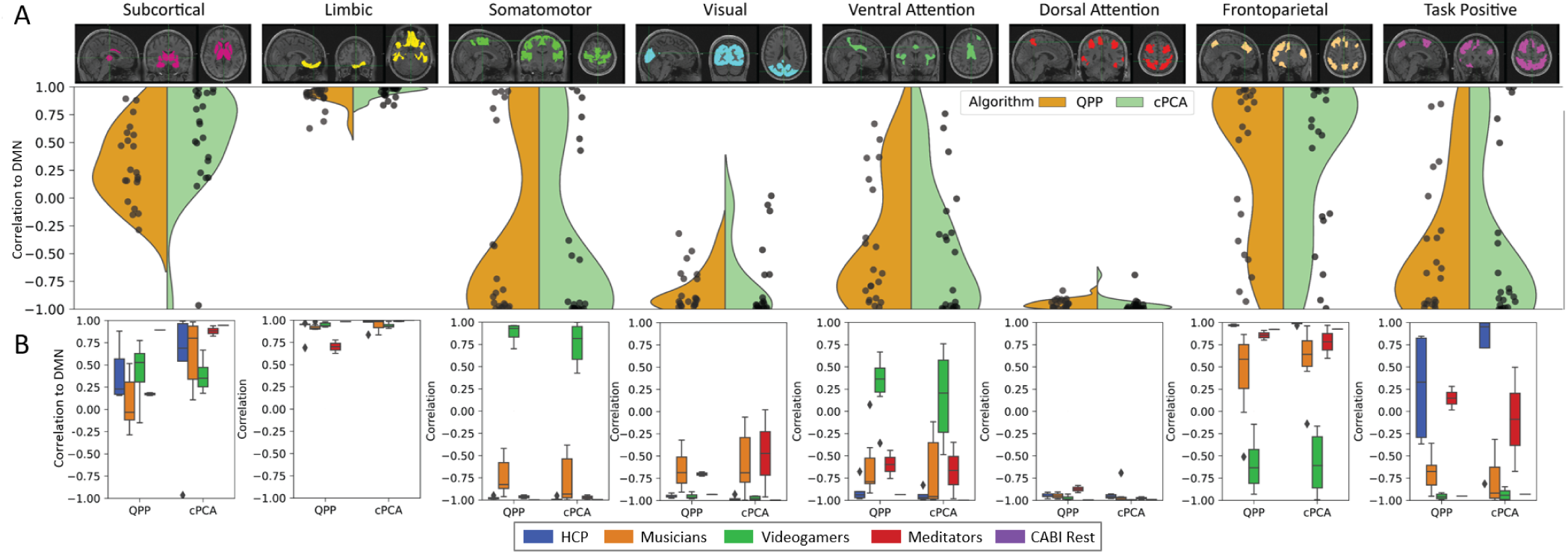
*Comparison of DMN Correlations Across Resampled TRs for QPP and cPCA Analyses.* (A) Violin plots displaying the distribution of DMN correlations across all datasets for both QPPs (yellow) and cPCA (green). Each network is displayed separately, with the corresponding brain regions highlighted above. (B) Boxplots showing the same DMN correlation distributions but separated by dataset, including HCP, Music, VG (Videogamers), Med (Meditators), and CABI Rest. Statistical analysis revealed no significant differences in DMN correlations between QPPs and cPCA across any dataset. A paired t-test between alogithm results confirmed this, as all datasets exceeded the significance threshold of 0.05: HCP (p=0.84), Music (p=0.56), VG (p=0.81), Med (p=0.74), and CABI (p=0.99). These results suggest that despite methodological differences, both QPP and cPCA approaches yield comparable DMN correlation patterns across datasets.

**Figure 7:**
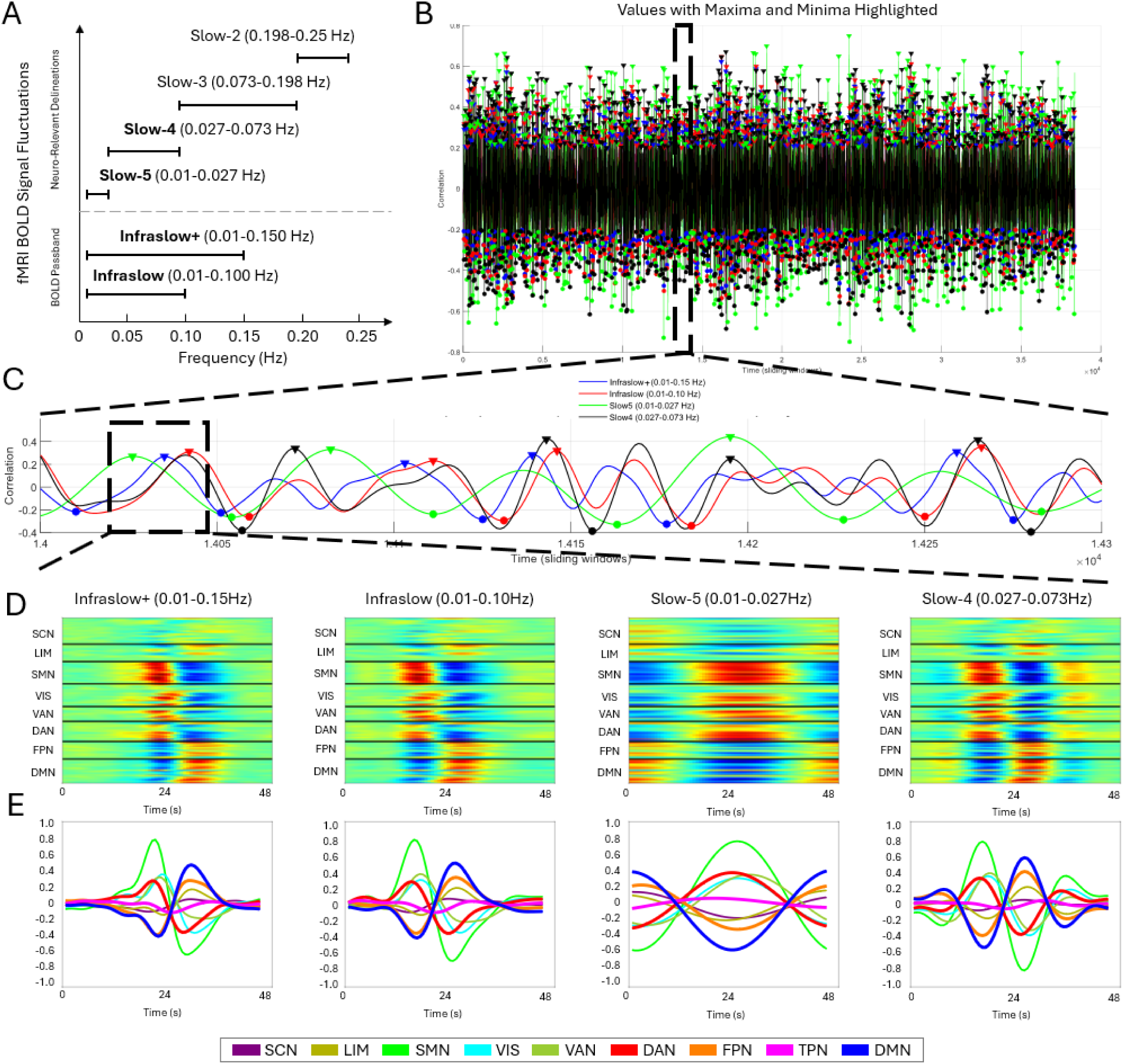
*Impact of Frequency Band Selection on QPP Detection on HCP Dataset.* (A) Illustration of the selected frequency bands and their relative placement within the fMRI BOLD signal fluctuation spectrum. (B) Full-group correlation of the four QPP templates, with maxima and minima highlighted to show distinct oscillatory patterns across frequency bands. (C) A zoomed-in segment of the QPP templates illustrating temporal shifts in correlation across networks. (D-E) QPP-derived templates for (D) Brainnetome and (E) Yeo with each frequency band, highlighting distinct network dynamics. Notably, Slow-5 emphasizes peak correlations between networks, while Slow-4, Infraslow, and Infraslow+ bands primarily capture spontaneous fluctuations and network transitions.

To quantitatively compare the fidelity of QPP and cPCA across TR variations, Figure 6 presents a detailed analysis of DMN correlations across all datasets. Violin and boxplots illustrate the distributions of DMN correlation values for both QPP and cPCA across resampled datasets. Statistical analysis revealed no significant differences in DMN correlations between QPP and cPCA within any dataset, as all paired *t*-test *p*-values exceeded the significance threshold of 0.05 (HCP: *p* = 0.84, Music: *p* = 0.56, Videogamers: *p* = 0.81, Meditators: *p* = 0.74, CABI: *p* = 0.99; see Appendix Table A3 for full results). At the same time, the distributions of DMN correlations vary across TR conditions and datasets, indicating that the absolute strength and dispersion of DMN coupling are influenced by temporal sampling and total scan length. Together, these findings suggest that while QPP and cPCA provide broadly similar DMN connectivity estimates, those estimates are modulated by TR and scan duration: fundamental network relationships are preserved across methods, but their precise magnitudes are sensitive to acquisition and resampling parameters.

Across datasets, the limbic network (LIM) consistently exhibited strong positive correlations with the DMN, reinforcing its intrinsic connectivity with core DMN regions. While the subcortical network showed some variation across datasets, it remained positively correlated with the DMN in a similar manner. In the sensory cortex, both the somatomotor (SMN) and visual (VIS) networks displayed strong anticorrelation with the DMN, except in the Videogamers dataset, where a more positive correlation was observed. Regarding the association cortex, the dorsal attention network (DAN) exhibited the strongest anticorrelation with the DMN across datasets, with the ventral attention network (VAN) also showing consistent anticorrelation. The frontoparietal network (FPN), however, demonstrated greater variability in its relationship with the DMN. Notably, the Videogamers dataset diverged from the other datasets, showing distinct correlation patterns with the DMN. Unlike other visual tasks where the somatomotor and visual networks were strongly anticorrelated with the DMN, the Videogamers dataset showed a more positive correlation in these networks, particularly in the VAN and SMN networks. This aligns with prior findings that externally cued visual tasks can exhibit task-negative-like responses in networks traditionally considered part of the task-positive network spectrum (Watters et al., 2025). Moreover, comparisons between Videogamers and other tasks revealed that the Videogamers task differed significantly from rest for all networks, though it showed only modest differences from the visual working memory task in the VAN response (Watters et al., 2025). These findings suggest that the Videogamers dataset represents a unique interaction between task engagement and intrinsic network dynamics, reinforcing the idea that dynamic shifts in DMN connectivity track a continuum between internal and externally driven attention states (Watters et al., 2025). Additionally, the task-positive network (TPN) exhibited inconsistent correlation with the DMN across datasets, likely reflecting that TPN can be divided into subnetworks with distinct relationships to the DMN.

Overall, these findings indicate that network dynamics remain preserved across datasets and TR conditions, reinforcing the validity of analyzing fMRI data acquired at slower sampling rates. Despite differences in scan duration, both QPP and cPCA successfully extract stable spatiotemporal patterns, suggesting that these approaches are effective for both individual-level and group-level analyses, even when lower TR-sampled data is compared to higher temporal resolution datasets.

### 3.3 Objective 3: Impact of Frequency Band Selection

Frequency band selection played a critical role in shaping QPP-derived and cPCA-derived network correlations, revealing distinct functional dynamics across different bands. Figure 7 illustrates the comparative effects of different frequency bands on QPPs and cPCA templates. Panel A outlines the frequency band definitions and their relative placement within the fMRI BOLD signal fluctuation spectrum. Across all frequency bands, QPP and cPCA analyses exhibited preserved network structures; however, distinct differences emerged in how each band captured temporal fluctuations in inter-network relationships.

A notable distinction was observed in the QPP-derived templates filtered within the Slow-5 band (0.01–0.027 Hz), which emphasized periods of maximal network synchronization—points at which inter-network correlations either peaked or reached their lowest values (Figure 7). This contrasts with results from QPPs filtered in the Slow-4 (0.027–0.073 Hz), Infraslow (0.01–0.10 Hz), and Infraslow+ (0.01–0.15 Hz) ranges, which instead captured transition points in network dynamics—reflecting shifts in spontaneous activity rather than stable periods of synchronization. Interestingly, the behavior of QPPs in the Slow-5 band closely resembled patterns typically captured by cPCA, as both emphasized sustained inter-network correlation structure, while QPPs in the Slow-4 band appeared more sensitive to transitions between these states. Notably, Slow-5 (0.01–0.027 Hz) is the only band that does not include the 0.05 Hz component, which has been identified as the dominant frequency for QPPs (Thompson et al., 2014). This suggests that Slow-5 filtering may isolate a distinct aspect of brain dynamics, potentially highlighting more stable inter-network coupling rather than the propagating quasi-periodic oscillations typically captured by QPPs at higher frequencies.

To further investigate the timing of network correlation shifts, we analyzed maxima (TMX1) and minima (TMX2) instances across frequency bands, as summarized in Table 5. The Wilcoxon signed-rank test revealed no significant difference between TMX1 and TMX2 instances (*p* = 0.21875), indicating that maxima and minima occurrences were evenly distributed across bands. These findings were further supported by an analysis of QPP detection stability across the scan (Appendix Figure A6). Across frequency bands, detected QPP TMX1 and TMX2 were uniformly distributed throughout the scan, suggesting that the presence of QPPs is not confined to specific time intervals. The comparable proportions of TMX1 and TMX2 further indicate that network fluctuations maintain a consistent balance between high and low correlation states, reinforcing the robustness of QPP detection regardless of the selected frequency band. However, Dunn’s test for post-hoc pair-wise comparison showed that Slow-5 exhibited a significant difference compared to other bands (*p* = 0.00216), suggesting that Slow-5 follows a distinct temporal pattern relative to other bands.

**Table 5:** Comparison of TMX1 (Maxima) and TMX2 (Minima) instances for each frequency band (left) and the Modified Jaccard Index for Maxima (TMX1) and Minima (TMX2) with Cushion (±8 timepoints) (right). Statistical analysis using the Wilcoxon signed-rank test revealed no significant difference between TMX1 and TMX2 (*p* = 0.21875). However, the Kruskal-Wallis test indicated a significant difference between Slow-5 and other frequency bands (*p* = 0.00216), suggesting that Slow-5 exhibits distinct spatiotemporal properties compared to the other bands. The **bold\*** values indicate statistically significant differences between groups using pair-wise comparison. All groups containing Slow-5 were not cyclically aligned with the other frequencies.

| Band | TMX1 (Maxima) | TMX2 (Minima) | Comparison | TMX1 (Maxima) | TMX2 (Minima) |
| --- | --- | --- | --- | --- | --- |
| Infraslow+ | 467 | 441 | Infraslow+ and Infraslow | 0.4311 | 0.4359 |
| Infraslow | 506 | 472 | Infraslow+ and Slow-5 | **0.0825\*** | **0.0807\*** |
| Slow-5 | 384 | 365 | Infraslow+ and Slow-4 | 0.3063 | 0.3063 |
| Slow-4 | 581 | 571 | Infraslow and Slow-5 | **0.0282\*** | **0.0227\*** |
|  |  |  | Infraslow and Slow-4 | 0.3534 | 0.3481 |
|  |  |  | Slow-5 and Slow-4 | **0.1058\*** | **0.0915\*** |

To quantify these differences, we measured the mean time lag between Slow-5 and the other frequency bands for both TMX1 and TMX2. The Slow-5 to Infraslow+ comparison resulted in a mean time difference of 4.53 timepoints (TMX1) and 4.92 timepoints (TMX2), while the Slow-5 to Infraslow comparison yielded a mean difference of 5.24 timepoints (TMX1) and 5.00 timepoints (TMX2). Finally, when comparing Slow-5 to Slow-4, the mean time difference was 4.04 timepoints (TMX1) and 3.94 timepoints (TMX2). An additional analysis with a cushion of 16 timepoints confirmed these findings, yielding the same statistical outcomes (Appendix Table A4). Because these lags are computed after aligning templates to maximize their cross-correlation, they should be interpreted as a relative shift in where the dominant component places the peak correlation/anticorrelation and major change-points in network activity, rather than as a literal onset delay in the underlying signal. This pattern reinforces the idea that Slow-5 captures a distinct oscillatory mode of the underlying signal compared to the other frequency bands. In contrast, Infraslow and Infraslow+ offered no significant difference in comparison, with the additional higher frequencies in Infraslow+ contributing noise but not meaningful distinctions in network correlation patterns.

The results indicate that Slow-5 captures temporally distinct patterns that coincide with periods of peak inter-network correlation, whereas templates in the other bands more frequently align with transitional segments of the trajectory (e.g., rising or falling phases of network coupling). This characteristic aligns closely with cPCA-derived templates (see Figure 8), which generally emphasize moments of maximal synchronization across networks. Across Infraslow, Slow-5, and Infraslow+ filtering, the leading cPCA component shows highly similar spatial structure, with the Slow-4 band exhibiting the largest deviation: its template is slightly shifted in phase and places somewhat more weight on intermediate network states, while preserving the same overall pattern of DMN–attentional and sensory interactions. For full cPCA components and explained variance, please see Appendix Figure A8. Notably, the significant temporal differences between Slow-5 and the other bands indicate that the Slow-5 component places its peaks of DMN–network correlation at a distinct phase of the trajectory, rather than simply reproducing the same change-points seen in Slow-4, Infraslow, and Infraslow+. Thus, while all bands reflect coordinated large-scale fluctuations, Slow-5 preferentially emphasizes epochs of maximal inter-network synchronization, consistent with the idea that this frequency range captures a particularly coherent mode of intrinsic connectivity.

**Figure 8:**
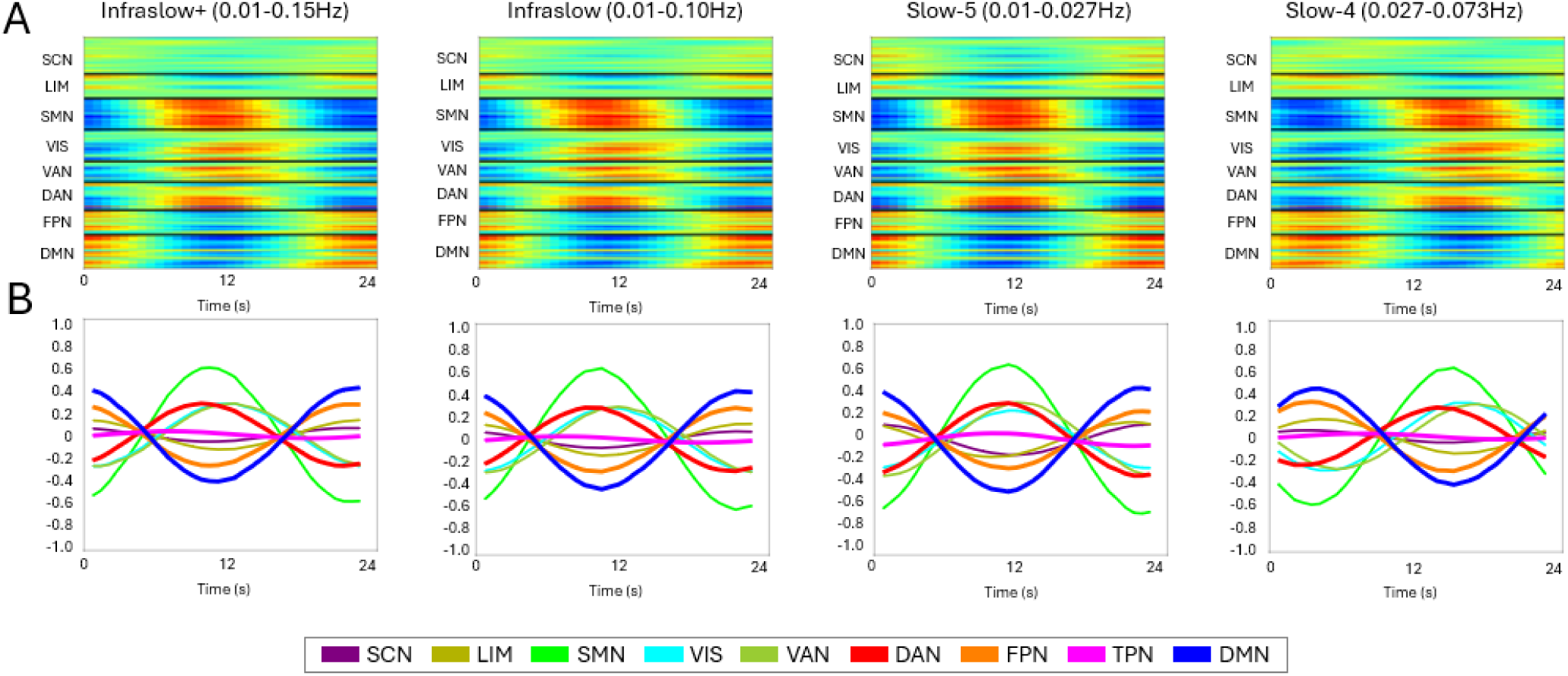
*Frequency Band Selection on cPCA Detection.* cPCA-derived templates for (A) Brainnetome and (B) Yeo with each frequency band. Infraslow, Slow-5, and Infraslow+ filtering yield highly similar leading components, emphasizing stable DMN–attentional and sensory network structure, whereas the Slow-4 band shows the largest deviation, with a modest phase shift while preserving the overall pattern.

Additionally, the Slow-5 window length analysis provided further insights into the temporal structure of detected QPPs. As illustrated in Appendix Figure A7, varying the window length within the Slow-5 band influenced the observed QPP structure in a manner that depended on its relationship to the dominant Slow-5 cycle. In this dataset, the primary Slow-5 component fluctuates within the 0.01–0.027 Hz band, corresponding to cycle lengths of approximately 37–100 s. When we systematically varied the Slow-5 window length, shorter windows (20–24 s) sampled only a portion of a Slow-5 cycle and tended to yield templates that emphasized periods of strong inter-network synchronization. As the window was extended (e.g., 32 s, 36 s, and beyond), the resulting QPPs increasingly incorporated segments where network relationships were changing, producing templates that more clearly reflected gradual transitions in network organization.

Taken together, these findings emphasize the importance of careful frequency band selection in functional neuroimaging studies, as different bands may highlight fundamentally different aspects of network dynamics. While Slow-5 provide a view of highly synchronized states, lower-frequency bands such as Slow-4 and Infraslow capture more gradual transitions in connectivity states. Furthermore, within Slow-5, the selection of window length plays a crucial role in determining whether QPP detection emphasizes short-term synchronization or longer-term network shifts. These distinctions have important implications for both individual and group-level analyses, particularly in studies examining dynamic brain states over varying temporal scales.

## 4 Discussion

The results demonstrate that both QPP and cPCA reliably identify large-scale spatiotemporal patterns across multiple datasets with different TRs and scan lengths, with both methods converging on similar core templates while still showing systematic modulation by acquisition choices. Despite differences in acquisition parameters, both approaches consistently captured resting-state network interactions, but they differed in their sensitivity to TR, scan duration, and inter-subject variability: QPP-derived templates were comparatively stable across resampling conditions, whereas cPCA-derived components showed larger changes in their network correlation profiles and DMN relationships. By systematically evaluating the impact of scan length, TR, and frequency band selection, this study provides critical insights into the methodological considerations that influence the detection and interpretation of dynamic functional connectivity patterns.

### 4.1 Impact of Scan Length on Stability and Variability in Dynamic Functional Connectivity

Prior studies have demonstrated that functional connectivity estimates become more reliable with increasing scan length (Birn et al., 2013; Laumann et al., 2015). Longer scans provide more occurrences of QPP cycles, allowing for better estimation of recurrent network dynamics, whereas shorter scans introduce greater variability and reduced reproducibility. Similar behavior can be expected for spatiotemporal patterns, and this challenge is particularly pronounced at the individual level. Here, we use the term *effective data duration* to refer to the total sampled time per subject (number of volumes multiplied by TR, summed across runs when applicable). When effective data duration is low—for example, due to the combination of short scans and long TRs—fewer QPP and cPCA cycles are observed, leading to higher variability in pattern detection. Stable pattern extraction in association and sensory networks may therefore require longer effective data durations. Subcortical networks, which inherently exhibit lower signal-to-noise ratios, also demonstrate greater instability under these conditions, further complicating individual-level pattern detection. Consequently, studies with limited effective data duration may need to aggregate data across multiple subjects to achieve stable and interpretable QPP and cPCA estimates (Gratton et al., 2018).

QPPs demonstrated greater reliability in capturing network dynamics at the individual level and exhibited increased stability with longer group scan durations, making them more suitable for detecting subject-specific differences in functional dynamics. In contrast, cPCA stabilized more quickly, offering computational efficiency for shorter scans but at the cost of higher individual-level variability, particularly in subcortical regions. This variability limits its feasibility for single-subject applications. However, QPPs remained effective in detecting subcortical patterns even at scan lengths feasible for individual analyses, suggesting they provide more consistent representations of network activity at this level.

Prior research suggests that approximately 20 minutes of scan time is sufficient to achieve reasonably stable group-level functional connectivity estimates in static analyses (Donnelly-Kehoe et al., 2019). Our findings are broadly consistent with this view, reinforcing the notion that longer scan durations improve the stability and reliability of QPP and cPCA detection, particularly for structured network dynamics. Studies using the HCP dataset have similarly reported increased reliability with longer scan times, with notable improvements observed in scans up to 30 minutes (Mejia et al., 2016). These results highlight the importance of acquiring sufficient scan lengths for robust functional connectivity analyses, especially in individual-level or clinical applications. While prior work points to an optimal range of approximately 20–30 minutes for stable group-level static connectivity (Donnelly-Kehoe et al., 2019; Mejia et al., 2016), these estimates are not directly optimized for dynamic spatiotemporal methods. Given that QPP and cPCA must capture recurring spatiotemporal patterns rather than time-averaged connectivity, our results suggest that dynamic analyses benefit at least as much from increased scan duration and may require more data to reach comparable levels of reliability, although we did not formally determine a minimum threshold. Although some studies suggest that incorporating spatial regularization can improve reliability even with shorter scan times (Duda et al., 2023), such techniques may also enhance QPP detection by providing a structured framework for identifying canonical patterns, potentially allowing for more efficient subject-specific analyses in future applications.

We observed network-specific differences in scan length requirements across both QPP and cPCA analyses. Association and sensory cortices exhibited high consistency across datasets and algorithms, reinforcing their robustness in spatiotemporal pattern characterization. In contrast, the frontoparietal network displayed increased variability at shorter scan durations, suggesting that longer scans are necessary for reliable detection in these regions. Subcortical networks exhibited the highest variability, particularly in cPCA estimates, but remained detectable with QPPs even at scan lengths feasible for individual-level analyses. This heightened variability in subcortical and limbic estimates is consistent with previous findings, such as those reported by Yousefi et al. (2021), where these regions showed lower BOLD amplitude and higher noise levels compared to cortical networks (Yousefi & Keilholz, 2021), and with work demonstrating that basic signal properties and complexity metrics vary systematically across networks and are strongly shaped by location-dependent signal properties (S. Keilholz et al., 2020). These factors likely reflect a combination of coil sensitivity profiles, vascular and physiological differences, and increased susceptibility to noise in deeper and medial structures. As a result, reliable extraction of subcortical QPP or cPCA patterns may require longer scan durations, targeted denoising strategies, or acquisition schemes that improve subcortical signal-to-noise ratio to achieve comparable fidelity to cortical networks.

These findings support previous research demonstrating that functional connectivity estimates become more stable with increasing scan time (Birn et al., 2013; Laumann et al., 2015). However, practical constraints—such as scanner availability, participant fatigue, and feasibility—often limit scan durations, particularly in clinical settings. In such cases, our results suggest that group-level aggregation or alternative modeling techniques (e.g., adaptive filtering, machine learning corrections) may be necessary to mitigate variability and optimize shorter scan analyses while maintaining analytical rigor (Gratton et al., 2018). Additionally, although individual-level analyses generally require longer scan durations, multivariate statistical techniques applied to large datasets with high subject numbers can compensate for shorter individual scans by leveraging population-level regularities. These methods may improve the reliability of functional connectivity estimates in studies with constrained scan times, an area warranting further investigation as new statistical approaches emerge.

### 4.2 Effects of Repetition Time on QPP and cPCA Sensitivity in Functional Brain Networks

TR is a critical factor in rs-fMRI studies, as it determines the temporal resolution and sampling density of the acquired data. Short TRs (e.g., *<* 1 s) can improve sensitivity to transient network dynamics, enabling more precise tracking of QPP propagation and phase-coherent cPCA structures by providing more samples per unit time (Belloy et al., 2018; Thompson et al., 2014). Conversely, longer TRs (*>* 2 s) may introduce undersampling artifacts, leading to temporal aliasing of faster vascular and neural fluctuations within the infraslow band, which can distort the apparent timing and waveform shape of these propagating patterns. Importantly, TR effects are not independent of scan duration. In shorter scans, longer TRs further reduce the effective number of QPP and cPCA cycles sampled, decreasing stability, whereas in longer scans, the increased total sampling time can partially mitigate these effects. Prior work has shown that TR influences functional connectivity estimates (Birn et al., 2013; Laumann et al., 2015), and these influences are expected to be even more pronounced in dynamic, propagating network patterns such as QPPs and cPCA-derived fluctuations.

While our findings suggest that QPP and cPCA remain largely robust across varying TRs in terms of large-scale network relationships, it is important to note that our analysis primarily focused on overall network correlation patterns rather than specific spatiotemporal features such as propagation speed and waveform structure. Prior studies indicate that higher temporal resolution is critical for capturing the precise timing of quasi-periodic wave propagation (Belloy et al., 2018; Thompson et al., 2014), and it remains possible that longer TRs introduce distortions in propagation trajectories, phase alignment, or oscillatory timing that were not directly assessed in our study. Future work should investigate how TR-dependent changes affect the fidelity of propagating network patterns, particularly in the context of disease-related alterations where timing disruptions may be diagnostically relevant.

Importantly, TR effects interact with scan duration because shorter TRs increase sampling density but reduce the signal available in each image, whereas longer TRs provide higher per-volume signal but require proportionally longer total scan time to achieve comparable effective data duration and reliability of dynamic estimates. This underscores the need for careful TR selection in study designs, particularly when comparing datasets acquired under different imaging conditions. Future studies should explore adaptive TR strategies, multi-resolution analyses, and advanced reconstruction techniques to optimize spatiotemporal pattern detection. Developing these methods will be essential for ensuring reproducibility across diverse imaging protocols and enhancing the applicability of QPP- and cPCA-based analyses to clinical and neuropsychiatric research.

### 4.3 Frequency Driven Variability in QPP and cPCA Across Network Dynamics

The choice of bandpass filter directly affects the detected frequency components of spatiotemporal patterns. Given the established role of infraslow fluctuations (0.01–0.1 Hz) in functional connectivity, different bandpass filters may emphasize distinct aspects of QPP and cPCA dynamics. Slow-5 (0.01–0.027 Hz) has been associated with physiological fluctuations, particularly in arterial partial pressure of CO_2_ and O_2_ (Lv et al., 2018), as well as metabolic and vascular processes (Gohel & Biswal, 2015). Given its lower frequency range, it may emphasize global network coherence rather than the propagating wave-like structure observed in broader infraslow bands that include the 0.05 Hz component. Consequently, while Slow-5 may capture aspects of long-range synchronization, it is important to interpret these findings cautiously, as it may not fully reflect the cyclic dynamics typical of QPPs. In contrast, Slow-4 (0.027–0.073 Hz) captures a broader spectrum of spontaneous neural fluctuations, including both localized and distributed activity. Prior studies have demonstrated that filtering choices alter functional connectivity networks (Zalesky et al., 2014), and given that QPP detection relies on waveform recurrence within specific frequency bands, it may be more sensitive to filtering choices than cPCA.

Our results indicate that frequency band selection significantly influences the stability and interpretability of QPP and cPCA-derived patterns. While Slow-4 (0.027–0.073 Hz), Infraslow, and Infraslow+ bands exhibited a high degree of homogeneity, capturing similar network dynamics, Slow-5 (0.01–0.027 Hz) showed distinct differences. Notably, when the window length (*WL*) was increased, Slow-5 led to a delayed timing of detected change points, suggesting that this band may highlight slower transitions in whole-brain activity compared to other infraslow bands. This delay may arise because Slow-5 predominantly reflects metabolic and vascular influences, which operate on slower time scales than neuronal oscillations. Unlike Slow-4, which includes more pronounced neural dynamics, Slow-5 may emphasize shifts in hemodynamic states that evolve over longer timescales. This could account for the timing shifts observed with increasing *WL* luctuations.

It is important to note that default preprocessing pipelines, such as the default CPAC bandpass filter (typically 0.01–0.1 Hz), may influence dynamic connectivity analyses by determining which frequency components are retained. Users should be aware that standard bandpass filtering choices may affect QPP and cPCA-derived patterns, particularly in analyses sensitive to frequency-dependent effects. The observed differences in Slow-5 versus Slow-4 highlight the need for careful consideration of band selection when interpreting dynamic functional connectivity results.

Despite these differences, QPPs were consistently detected across all frequency bands, underscoring their robustness in tracking spontaneous whole-brain activity. The stability of QPPs across Slow-4, Infraslow, and Infraslow+ bands suggests that they remain a reliable marker of large-scale network dynamics, even amid preprocessing variations. However, in Slow-5, increasing *WL* shifted the timing of change points, indicating that this band may influence the temporal resolution of detected transitions in functional states. In contrast, cPCA remained largely stable across frequency bands but exhibited subtle differences in network synchronization patterns, particularly in subcortical and frontoparietal regions. This distinction suggests that while QPPs are effective in detecting transient reorganization of network states, cPCA may provide a more stable representation of underlying functional architecture. Notably, in frequency bands such as Slow-5, where quasi-periodic dynamics may be less pronounced, cPCA’s emphasis on stationary network features could complement QPP-based analyses by identifying consistent background connectivity patterns. Future work could extend these findings by examining even higher-frequency bands, where network modularity and fine-grained oscillatory dynamics may play a more prominent role in shaping functional connectivity patterns (DeRamus et al., 2021).

A key open question is whether cross-frequency interactions influence QPP dynamics. Given that functional connectivity fluctuations emerge from nested oscillations across frequency bands, Slow-5, in particular, may serve as a bridge between hemodynamic and neuronal oscillations, reflecting vascular contributions that modulate faster neural activity. This aligns with previous findings that suggest low-frequency fluctuations act as a carrier signal for higher-frequency cortical oscillations, influencing network dynamics at multiple timescales (Hutchison et al., 2013). Furthermore, QPPs are modulated by external sensory input, as Belloy et al. (2018) demonstrated that infraslow quasi-periodic patterns respond to visual stimuli (Belloy et al., 2018), and Xu et al. (2023) highlighted how large-scale network interactions adapt to sensory inputs (Xu, Smith, et al., 2023). Understanding these cross-frequency and sensory interactions may provide deeper insights into the hierarchical organization of brain rhythms.

Although Slow-4 effectively captures infraslow dynamics, our results suggest that Slow-5 may contain additional signals relevant to long-range synchronization effects or propagation characteristics in brain networks. Given these distinctions, future research should consider multi-band approaches, integrating insights from both Slow-4 and Slow-5 to develop a more comprehensive understanding of QPP-based connectivity dynamics.

### 4.4 The Role of Spatiotemporal Patterns in Functional Connectivity Analysis

Yousefi et al. (2018) demonstrated that QPPs are prominent features of intrinsic brain activity that contribute substantially to functional connectivity, with networks such as the DMN and task-positive systems emerging as major contributors in HCP data (Van Essen, Smith, et al., 2013; Yousefi et al., 2018). Abbas et al. (2019) further showed that QPP spatiotemporal patterns differ between rest and stimulation, strongly modulating DMN–TPN interactions and highlighting QPPs as key mechanisms shaping large-scale connectivity (Abbas et al., 2019).

Maltbie et al. (2022) compared several dynamic functional connectivity methods—including sliding-window correlation (SWC), co-activation patterns (CAPs), and phase-synchrony clustering—and found that CAP-based approaches are particularly sensitive to QPP sequences (Maltbie et al., 2022). CAPs and related state-based methods tend to segment different phases of the QPP cycle into discrete network states, whereas SWC treats QPPs as transient fluctuations within a slowly varying connectivity profile. Thus, QPPs interact non-trivially with state-clustering approaches and their phase progression likely corresponds to transitions between data-driven connectivity states.

Complementing this view, Bolt et al. (2022) proposed a parsimonious framework in which large-scale brain activity is described as a combination of standing waves (zero-lag synchrony) and traveling waves (time-lag synchrony), using cPCA to bridge conventional connectivity measures with QPP-like propagations (Bolt et al., 2022). This work reinforces the idea that many dynamic methods capture overlapping components of a common spatiotemporal structure, with QPPs emphasizing propagating dynamics and cPCA providing a unified representation of both synchronous and time-lagged activity.

QPP-driven DMN activity, associated with internally oriented cognition such as mind-wandering and self-referential thought (Raichle et al., 2001), has been linked to attention shifts and creative ideation (Seeburger et al., 2024; Watters, Fazili, Daley, Belden, LaGrow, et al., 2024; Watters et al., 2025), underscoring the behavioral relevance of these propagating patterns. Altogether, the spatiotemporal templates extracted by the QPP algorithm and cPCA represent biologically meaningful constructs that summarize large-scale neural dynamics rather than purely statistical artifacts.

By directly evaluating QPPs and cPCA across diverse datasets and systematically varying scan duration, TR, and frequency band selection, the present study provides an empirical basis for using these methods in dynamic functional connectivity research. Our results extend prior work by testing their reproducibility and sensitivity under acquisition and preprocessing conditions that mirror real-world neuroimaging protocols. In doing so, we show that QPP and cPCA converge on similar large-scale spatiotemporal patterns while offering complementary strengths—QPPs provide stable, interpretable templates even at scan lengths relevant for individual-level inference, whereas cPCA offers a broader view of phase-coherent fluctuations—supporting their use as practical tools for studying intrinsic brain activity in both basic and clinical neuroscience.

### 4.5 Methodological Considerations and Variability in QPP Detection

One important consideration is the impact of atlas selection on spatiotemporal pattern detection. While this study used the Brainnetome ROIs and Yeo Resting State Networks (RSNs), other parcellations—such as Schaefer (Schaefer et al., 2018), AAL (Automated Anatomical Labeling) (Tzourio-Mazoyer et al., 2002), and NeuroMark (Iraji et al., 2023; Jensen et al., 2024)—may produce different QPP and cPCA templates due to variations in spatial resolution and network definitions. Previous studies have demonstrated that different brain parcellations yield systematically different functional connectivity estimates (Zalesky et al., 2014), raising concerns about methodological standardization. Although these atlases vary in their granularity and methodological assumptions, we selected the Yeo and Brainnetome parcellations for their consistency with prior studies and their widespread use in functional connectivity research. The Yeo

RSNs provide a widely adopted framework, making them suitable for large-scale connectivity analysis, while the Brainnetome ROIs offer a finer-grained parcellation that better captures regional specificity. By using these two complementary parcellations, our study ensures comparability with previous findings while maintaining network-level and regionally detailed insights into spatiotemporal dynamics.

Another key issue in spatiotemporal pattern detection is the role of global signal regression (GSR), a widely debated preprocessing step in resting-state fMRI analysis. GSR removes the global mean time series from voxel-wise signals to reduce widespread fluctuations that can reflect both neural activity and non-neural confounds. Although GSR can improve the apparent specificity of functional connectivity by attenuating physiological and motion-related artifacts (Murphy et al., 2009; Power et al., 2014), it remains controversial because it can induce anti-correlations and alter the covariance structure of the data (T. T. Liu et al., 2017; Murphy & Fox, 2017). In our data, both QPPs and cPCA robustly recover global spatiotemporal structure, including components that persist after GSR, consistent with the idea that QPPs reflect a structured contribution to the global signal rather than being purely noise-driven. To address these concerns, the authors also repeated the full analysis without GSR (see supplemental analyses in Watters et al. (2025)); as expected, this yielded strong global correlations and did not produce a prominent DMN anti-correlation QPP, indicating that the principal spatiotemporal structure is not an artifact of the regression step.

In this study, we applied GSR to focus on network dynamics beyond the global signal. As shown by Bolt et al., 2022, the first principal component extracted from both QPP and cPCA analyses closely corresponds to the global signal, confirming that global fluctuations dominate early components of spatiotemporal patterns. While the global signal contains meaningful neural information, our primary objective was to isolate and analyze structured network dynamics beyond these widespread global fluctuations. Applying GSR ensures that our findings emphasize regional network interactions rather than being driven by global signal variations, making the results more relevant for understanding localized and propagating connectivity changes across brain networks.

A critical methodological shift in dynamic functional connectivity analysis is moving beyond traditional terminology such as ”task-positive network” (TPN) when defining network-specific correlations. Historically, TPN has been used to contrast anti-correlated networks (e.g., DMN vs. TPN), but this definition primarily originated from seed-based analyses, which may oversimplify the complexity of whole-brain network interactions. Instead, studies increasingly emphasize more precise descriptions of network dynamics that account for overlapping and co-fluctuating regions across multiple states. Depending on the research question, a more nuanced approach to defining functional networks—rather than rigid task-positive/task-negative dichotomies—may be necessary to accurately interpret QPP-derived patterns and their relationship to cognitive states.

### 4.6 Limitations

A key strength of this study lies in its multi-dataset design and systematic TR and scan-length sampling, which enhance the generalizability of the findings. By incorporating datasets with diverse acquisition parameters and scan conditions, we were able to evaluate how TR, scan duration, and frequency band selection affect QPP and cPCA estimates under conditions that approximate real-world neuroimaging protocols.

Despite these strengths, several methodological limitations must be acknowledged. First, multi-dataset analyses are inherently influenced by differences in acquisition protocols and scanner hardware. To mitigate this, we standardized preprocessing across all datasets using the Configurable Pipeline for the Analysis of Connectomes (C-PAC) (Craddock et al., 2013), applying consistent motion correction, spatial normalization, and filtering. This reduces, but does not eliminate, residual variability arising from site-and protocol-specific factors, underscoring the need for continued harmonization efforts in large-scale neuroimaging work.

Second, differences in cognitive state across datasets can substantially shape the expression of spatiotemporal patterns. In particular, the Videogamers dataset was collected during an active task context rather than true rest (a moving-dot reaction time paradigm), and task engagement can bias or reorganize intrinsic dynamics relative to conventional resting-state acquisitions. Rather than undermining these approaches, this contrast reinforces that QPP and cPCA are sensitive to both spontaneous fluctuations and task-modulated activity, and that apparent discrepancies across datasets may reflect genuine state dependence. This interpretation is consistent with prior work showing that infraslow and global spatiotemporal structure can differ systematically between resting and task conditions (Watters et al., 2025). Future studies should more explicitly control and monitor behavioral state, for example, through behavioral logs, task regressors, and standardized rest–task transition designs, to better dissociate spontaneous neural activity from task-induced (and post-task) influences on QPP expression.

Finally, QPP and cPCA detection relies on BOLD fMRI fluctuations, which provide indirect and temporally smoothed indices of neural activity. While resting-state fMRI is well-suited for mapping large-scale dynamics, its limited temporal precision constrains inferences about fast electrophysiological processes. Multimodal approaches integrating EEG with fMRI have shown that QPPs in resting-state fMRI correlate with infraslow electrical activity (Grooms et al., 2017; Thompson et al., 2014), and studies of non-stationary BOLD connectivity fluctuations have linked dynamic fMRI changes to electrophysiological oscillations (Tagliazucchi et al., 2013). Nonetheless, additional work is needed to more precisely map the neural generators and frequency-specific contributions underlying the propagating patterns captured by QPP and cPCA.

#### 4.6.1 Scan Length Considerations

- **Longer scan durations improve reliability**: QPPs and cPCA become more reliable as more quasi-periodic cycles are sampled. In our analyses, datasets with only a few minutes of data per condition yielded detectable but more variable patterns, whereas reliability—defined here as the consistency of extracted QPP and cPCA patterns across repeated analyses and independent datasets—increased substantially once scan durations exceeded approximately 10 minutes. This is particularly relevant for individual-level analyses, where limited data can increase variability in QPP amplitude and phase. TR also plays a role: shorter TRs (*<* 1 s) provide more samples per cycle and therefore better resolve fine-grained features of the waveform (e.g., peak timing and rapid changes in inter-network coupling), whereas longer TRs (*>* 2 s) undersample these dynamics. When feasible, multi-session aggregation should be used to increase effective data duration and reduce session-specific variability. Interpolation or spatial–temporal regularization techniques may further help compensate for lower temporal sampling in longer-TR scans while preserving key QPP dynamics.
- **Shorter scans favor group-level aggregation**: For very short scans (on the order of a few minutes), variability increases markedly for individual-level estimates. Our findings suggest that group-level averaging or multi-session aggregation is often necessary to obtain reliable QPP and cPCA representations under these constraints. Additionally, leveraging precomputed QPP templates or applying structural constraints, as suggested in prior work (Duda et al., 2023), may help stabilize pattern estimation in limited data settings and guide future efforts toward more robust subject-specific QPP analyses.
- **Subcortical networks require special consideration**: Subcortical regions exhibited greater variability, particularly in cPCA estimates, likely due to lower signal-to-noise ratios and greater susceptibility to physiological and hemodynamic fluctuations. Longer scan durations and larger sample sizes improve the reliability of subcortical spatiotemporal patterns, but region-specific filtering, tailored denoising, or multimodal integration (e.g., EEG–fMRI) may also be necessary for robust extraction in these structures.

#### 4.6.2 Repetition Time (TR) Optimization

- **QPPs and cPCA remain broadly robust across TRs**: Within the range of TRs examined in our resampling analyses, core large-scale network relationships (e.g., interactions between DMN and attention networks) were preserved, even though fine-scale temporal details of the waveforms varied. This supports the use of QPP- and cPCA-based approaches in datasets acquired with moderately different TRs.
- **Interpret longer TRs with appropriate caution**: Longer TRs reduce temporal sampling density and may obscure rapid changes in waveform shape or propagation timing, particularly for quasi-periodic waves. Our results suggest that large-scale propagating patterns remain detectable at coarser TRs, but we did not directly evaluate clinical datasets such as ADNI or PPMI in this work. Consequently, applications to clinical protocols with TRs in the 2–3 s range should be accompanied by careful consideration of which aspects of the dynamics (e.g., coarse network relationships vs. precise propagation timing) are interpretable.
- **Adjust scan duration to compensate for TR**: Datasets with longer TRs should use correspondingly longer total scan times to achieve comparable effective data duration and reliability of dynamic estimates. Future research should explore adaptive TR strategies, multi-resolution analyses, and advanced reconstruction methods to optimize QPP and cPCA detection across diverse imaging protocols.

#### 4.6.3 Frequency Band Selection

QPPs exhibited broadly consistent spatial structure across the Slow-4, Infraslow, and Infraslow+ ranges, making them well-suited for detecting change points in network dynamics and spontaneous shifts in functional connectivity. In contrast, cPCA primarily captured phase-coherent synchronization of network correlations, providing complementary insight into coherent oscillatory relationships within intrinsic brain activity. Selecting an appropriate frequency range is therefore crucial for optimizing QPP and cPCA applications, although the most informative band will depend on both acquisition parameters and the specific physiological question.

- **Slow −5 emphasizes recurrent structure; Slow-4 highlights transitions**: In our data, Slow-5 (0.01–0.027 Hz) tended to emphasize long-range synchronization and recurrent QPP cycles, with shorter window lengths (approximately 20–30 s) often yielding templates that sampled segments of strong inter-network correlation. Slow-4 (0.027–0.073 Hz), on the other hand, more frequently produced templates that highlighted network transitions and dynamic connectivity shifts. These patterns suggest that Slow-5 may be particularly informative for studying propagating spatiotemporal waves and recurrent motifs, whereas Slow-4 may better capture change points in coupling; however, the relative advantages of these bands—and their dependence on window length and dataset—require further systematic investigation.
- **Consider multi-band QPP analysis rather than a single fixed band**: Distinct frequency ranges accentuate different facets of neural dynamics. In our analyses, Slow-5 tended to highlight highly synchronized oscillatory motifs, whereas Slow-4 captured broader shifts in network state and change-points in coupling. Exploring both bands side by side, rather than committing to a single range *a priori*, can therefore uncover complementary information. The most informative band will likely depend on the physiological process of interest; integrating rs-fMRI with modalities such as EEG or functional ultrasound may help identify which bands best capture relevant activity patterns and whether combining bands improves sensitivity to both recurrent and transient motifs. Because the neurophysiological specificity of Slow-5 is still not fully established, findings in this band should be interpreted cautiously and, when possible, validated in follow-up multimodal or longitudinal work.

By carefully selecting frequency ranges and leveraging multi-band analysis, researchers can obtain a more comprehensive view of spatiotemporal network dynamics, ensuring that both relatively stable and transient components of functional connectivity are captured.

## 5 Conclusion

In this study, we evaluated the effects of scan duration, repetition time (TR), and frequency band selection on the detection and stability of QPPs and cPCA in resting-state fMRI. Our findings provide methodological insights and practical recommendations for optimizing spatiotemporal pattern detection in neuroimaging research.

Longer scan durations improve stability, particularly for individual-level analyses, with QPPs showing higher reproducibility at shorter scan lengths and cPCA benefiting from group-level aggregation. Subcortical networks were more sensitive to scan duration, requiring longer acquisitions for reliable detection.

Both QPPs and cPCA remained robust across different TRs, preserving core network relationships such as the DMN–DAN anti-correlation, reinforcing their utility in clinical fMRI studies with longer TR constraints. Frequency band selection also influenced pattern stability: in our analyses, Slow-5 (0.01–0.027 Hz) tended to emphasize recurrent, highly synchronized network structure, whereas Slow-4 (0.027–0.073 Hz) more often highlighted dynamic connectivity shifts and transitions in coupling. Multi-band QPP extraction is therefore recommended to probe both recurrent motifs and state changes, while future work should further clarify how band choice interacts with acquisition parameters and disease-related effects.

QPPs and cPCA offer complementary strengths. Using both methods together enhances the characterization of functional organization. By refining methodological approaches and ensuring robust analytic strategies, we can enhance the reliability of spatiotemporal pattern analysis in resting-state fMRI and pave the way for novel applications in cognitive neuroscience, clinical diagnostics, and network neuroscience.

## Data and Code Availability

Matlab scripts for analysis and visualization (Xu, Yousefi, et al., 2023): https://github.com/imnanxu/ QPPLab. The cPCA method applied to BOLD signal is based on finding by (Bolt et al., 2022) and packaged into Python scripts for analysis and visualization, available on Github: https://github.com/ tsb46/complex pca. The MRI preprocessing code and pipeline are open-source and publicly available (Craddock et al., 2013): https://fcp-indi.github.io/. The analysis and visualization performed for this paper utilized Matlab (Inc., 2022), Python 3.9 (Van Rossum & Drake Jr, 1995), R (R Core Team, 2021), and Bash scripting for long processing. Analysis scripts for demonstration are available on the Keilholz MIND Lab’s Github: https://github.com/GT-EmoryMINDlab.

All original scan acquisition adhered to best practices for ethical human data collection and informed consent in accordance with local institutional review boards. See original data acquisition study for details: HCP: Van Essen, Smith, et al., 2013; Videogamers: Jordan and Dhamala, 2023; Musicians: Belden et al., 2020; Meditators: Hasenkamp et al., 2012; and CABI Rest: Godwin et al., 2017.

## Author Contributions

Conceptualization, ideation, organization, data preparation, data processing, formal analysis, figure drafting, paper writing, review, editing: TJL. Data preparation, preprocessing troubleshooting, review, and editing: HW. Ideation, review, and editing: LD, VI, DS, NA, AF, MAK. Supervision, review, and editing: VC, WJP, EHS, SK.

## Funding

Research reported in this publication was supported by 1R01AG062581. Additionally, this work was supported by Office of Naval Research grant N00014-22-1-2218 to Randall W. Engle. Further, TJL would like to thank the financial support from the George R. Riley Fellowship.

## Declaration of Competing Interests

The authors declare they have no competing interests.

## Acknowledgements

The authors would like to extend their deepest gratitude to the patients and clinicians whose time, effort, and participation made the creation of these datasets possible. Though their contributions may go unnamed and unseen, they form the foundation upon which this research, and the broader field of neuroscience, continues to advance. We thank them tremendously.

We also acknowledge and thank the original data contributors and investigators of the Human Connectome Project, the Videogamers dataset (Georgia State University), the Musicians dataset (Northeastern Biomedical Imaging Center and Olin Neuropsychiatry Research Center), the Meditators dataset (Emory University), and the CABI Rest dataset (Georgia Institute of Technology). Their commitment to open science and data sharing has made this work possible.

## Notes

### Competing Interest Statement

The authors have declared no competing interest.

