## Supplementary and Appendix for "Fidelity of Spatiotemporal Patterns of Brain Activity Across Sampling Rate, Scan Duration, and Frequency Content"

January 2, 2026

**Keywords:** Resting-state fMRI, Spatiotemporal patterns, Quasi-Periodic Patterns (QPPs), Complex Principal Component Analysis (cPCA), Functional connectivity, Neuroimaging methodology, Repetition time (TR), Frequency band selection, Scan duration, Intrinsic brain dynamics

### Supplementary Material

#### Dataset Imaging Parameters

| Dataset | Source | T1 Anatomical | Functional Scan |
|---|---|---|---|
| Human Connectome Project (HCP) | Human Connectome Project. Washington University. (Van Essen et al., 2013). | Siemens Skyra 3T; 3D MPRAGE sequence; TR = 2400 ms, TE = 2.14 ms, TI = 1000 ms, FA = 8°, FOV = $224 \times 224$ mm, voxel size = 0.7 mm$^3$. | Siemens Skyra 3T; Gradient-echo Echo Planar Imaging; TR = 720 ms, TE = 33.1 ms, FA = 52°, FOV = $208 \times 180$ mm (RO x PE), matrix = $104 \times 90$ (RO x PE), slice thickness = 2.0 mm; 72 slices; 2.0 mm isotropic voxels, multi-band factor = 8, echo spacing = 0.58 ms. |
| Videogamers | Dr. Mukeshwar Dhamala (Georgia State University) and Timothy Jordan (UCLA). Georgia State/Georgia Tech Center for Advanced Brain Imaging (CABI) (Jordan & Dhamala, 2023). | Siemens 3T Magnetom Prisma MRI scanner; TR = 2530 ms; TE1-4: 1.69–7.27 ms; flip angle = 7°; Voxel Size: 1 mm$^3$. | Siemens 3T Magnetom Prisma MRI scanner; TR = 535 ms; TE = 30 ms; flip angle = 46°; FOV = 240 mm; Voxel Size: $3.8 \times 3.8 \times 4$ mm; slices: 32. |
| Musicians | Dr. Alexander Belden and Dr. Psyche Loui, Northeastern Biomedical Imaging Center and the Olin Neuropsychiatry Research Center (Belden et al., 2020). | Siemens 3T scanner; 3D magnetization prepared rapid-acquisition gradient-echo (MPRAGE) sequence; voxel size = $0.8 \times 0.8 \times 0.8$ mm$^3$ (TR = 2400 ms, TE = 2.09 ms, flip angle = 8°, FOV = 256 mm). | Siemens 3T scanner; echo-planar imaging (EPI) sequence; 947 volumes; voxel size = $3.0 \times 3.0 \times 3.0$ mm$^3$ (TR = 475 ms, TE = 30 ms, flip angle = 90°, 48 slices, FOV = 240 mm). Participants were instructed to keep their eyes open and fixated on a cross for the duration of the 7.5-minute resting-state scan. |
| Meditators | Dr. Wendy Hasenkamp, Dr. Larry Barsalou, and Dr. Christine Wilson-Mendenhall, Emory University. (Hasenkamp et al., 2012). | Siemens 3T Trio MRI scanner; TR = 2600 ms; TE = 3.9 ms; FOV = 240 mm. | Siemens 3T Trio MRI scanner; TR = 1500 ms; TE = 30 ms; flip angle = 90°; FOV = 192 mm; Voxel Size: $4 \times 3 \times 3$ mm; slices: 18. |
| CABI Rest | Dr. Eric Schumacher, Georgia Institute of Technology. Georgia State/Georgia Tech Center for Advanced Brain Imaging (CABI). Godwin et al., 2017 | Siemens 3T Trio MRI scanner; TR = 2250 ms; TE = 3.98 ms; flip angle = 9°; FOV = 256 mm; Voxel Size: 1 mm$^3$. | Siemens 3T Trio MRI scanner; TR = 2000 ms; TE = 30 ms; flip angle = 90°; FOV = 240 mm; Voxel Size: $3 \times 3 \times 3$ mm; slices: 37. |

**Table S1:** Details of datasets, sources, and MRI scan parameters.

#### Brannetome ROIs Mapped to Yeo's Networks

| Network | ROI Indices |
|---|---|
| Default Mode Network (DMN) | 3, 5, 6, 11, 13, 14, 23, 33, 34, 35, 41, 42, 43 |
| | 44, 51, 52, 79, 80, 81, 83, 84, 87, 88, 95, 121, 122 |
| | 141, 144, 153, 154, 175, 176, 179, 181, 187, 188 |
| Ventral Attention Network (VAN) | 2, 15, 37, 38, 39, 40, 61, 62, 65, 123, 124, 167, 168 |
| | 169, 170, 173, 174, 180, 183, 184, 185, 186 |
| Dorsal Attention Network (DAN) | 7, 8, 25, 26, 30, 55, 56, 63, 64, 85, 86, 91, 92 |
| | 97, 98, 107, 125, 126, 127, 128, 129, 130, 133, 134, 139, 140 |
| | 143, 150, 159, 201 |
| Frontoparietal Network (FPN) | 1, 4, 12, 16, 17, 18, 19, 20, 21, 22, 24, 28, 29 |
| | 31, 32, 36, 46, 82, 99, 100, 137, 138, 142, 147, 148, 166 |
| Task-Positive Network (TPN) | 7, 8, 25, 26, 30, 55, 56, 63, 64, 85, 86, 91, 92 |
| | 97, 98, 107, 125, 126, 127, 128, 129, 130, 133, 134, 139, 140 |
| | 143, 150, 159, 201, 1, 4, 12, 16, 17, 18, 19, 20, 21, 22, 24 |
| | 28, 29, 31, 32, 36, 46, 82, 99, 100, 137, 138, 142, 147, 148, 166 |
| Visual Network (VIS) | 105, 106, 108, 112, 113, 114, 119, 120, 135, 136, 151, 152, 182 |
| | 189, 190, 191, 192, 193, 194, 195, 196, 197, 198, 199, 200, 202 |
| | 203, 204, 205, 206, 207, 208, 209, 210 |
| Somatomotor Network (SMN) | 9, 10, 53, 54, 57, 58, 59, 60, 66, 67, 68, 71, 72 |
| | 73, 74, 75, 76, 131, 132, 145, 146, 149, 155, 156, 157, 158 |
| | 160, 161, 162, 163, 164, 171, 172 |
| Limbic Network (LBN) | 27, 45, 47, 48, 49, 50, 69, 70, 77, 78, 89, 90, 93 |
| | 94, 96, 101, 102, 103, 104, 109, 110, 111, 115, 116, 117, 118 |
| Subcortical Network (SCN) | 165, 177, 178, 211, 212, 213, 214, 215, 216, 217, 218, 219, 220 |
| | 221, 222, 223, 224, 225, 226, 227, 228, 229, 230, 231, 232, 233 |
| | 234, 235, 236, 237, 238, 239, 240, 241, 242, 243, 244, 245, 246 |

**Table S2:** ROI indices associated with each functional network.

### 1 Appendix

#### 1.1 Global Signal Regression Representation

$$X' = X - (Xg)\frac{g^T}{g^T g}$$

where:

- $X$ is the $N \times T$ matrix of fMRI time series (with $N$ regions and $T$ time points),
- $g = \frac{1}{N}\sum_{i=1}^{N} X_i$ is the global signal (the mean time series across all regions),
- $Xg$ is the projection of $X$ onto $g$,
- $X'$ is the GSR-corrected time series.

#### 1.2 Modified Jaccard Index with Cushion

Let $S_1$ and $S_2$ be two sets of numerical values, and let $\tau$ be the cushion (tolerance for overlap). The overlap count is given by:

$$O_\tau = \sum_{s_1 \in S_1} 1\left(\min_{s_2 \in S_2} |s_1 - s_2| \le \tau\right)$$

where $1(\cdot)$ is the indicator function that returns 1 if the condition inside holds and 0 otherwise.

The union count is computed as:

$$U = |S_1 \cup S_2|$$

Thus, the Modified Jaccard Index with Cushion is given by:

$$J_\tau(S_1, S_2) = \frac{O_\tau}{U}$$

To assess whether the Jaccard indices differ significantly across conditions, a statistical analysis was conducted. Specifically, the Wilcoxon Signed-Rank Test was employed to compare Jaccard indices between maxima (TMX1) and minima (TMX2), given the non-normal distribution of the data.

Additionally, the Mann-Whitney U Test was used to evaluate whether specific Jaccard index pairs significantly differed from other frequency band pairings, with significance determined at $p < 0.05$. If significance is found, a precursor analysis is performed to evaluate whether cyclic activity exhibits coherence across conditions.

#### 1.3 Precursor Time Point Analysis

Let $T_{\max}^{(f_1)}$ and $T_{\min}^{(f_1)}$ be the time indices of maxima and minima in frequency band $f_1$, and let $T_{\max}^{(f_2)}$ and $T_{\min}^{(f_2)}$ be the corresponding time indices in frequency band $f_2$. The tolerance threshold for matching time points is denoted by $\tau$ (e.g., $\pm 8$ time points). The minimum absolute time differences for maxima and minima between the two frequency bands are given by $\delta_{\max}^{(f_1,f_2)}$ and $\delta_{\min}^{(f_1,f_2)}$, respectively.

The Precursor Analysis for Maxima is given by:

$$\delta_{\max}^{(f_1,f_2)} = \min_{t_2 \in T_{\max}^{(f_2)}} |t_1 - t_2|, \quad \forall t_1 \in T_{\max}^{(f_1)}$$

Similarly, the Precursor Analysis for Minima is:

$$\delta_{\min}^{(f_1,f_2)} = \min_{t_2 \in T_{\min}^{(f_2)}} |t_1 - t_2|, \quad \forall t_1 \in T_{\min}^{(f_1)}$$

To compute the Mean Time Differences for matched maxima and minima within the given tolerance $\tau$:

$$\bar{\delta}_{\max}^{(f_1,f_2)} = \frac{1}{|T_{\max,\tau}^{(f_1)}|} \sum_{t \in T_{\max}^{(f_1)}} \delta_{\max}^{(f_1,f_2)} 1(\delta_{\max}^{(f_1,f_2)} \leq \tau)$$

$$\bar{\delta}_{\min}^{(f_1,f_2)} = \frac{1}{|T_{\min,\tau}^{(f_1)}|} \sum_{t \in T_{\min}^{(f_1)}} \delta_{\min}^{(f_1,f_2)} 1(\delta_{\min}^{(f_1,f_2)} \leq \tau)$$

where $T_{\max,\tau}^{(f_1)}$ and $T_{\min,\tau}^{(f_1)}$ are the subsets of time points within the tolerance $\tau$.

This method allows a direct comparison of how close extrema (maxima/minima) are across frequency bands.

#### 1.4 (Objective 1) Demonstration of Linear X-Axis for Scan Length Individual Feasibility vs. Group-Analysis

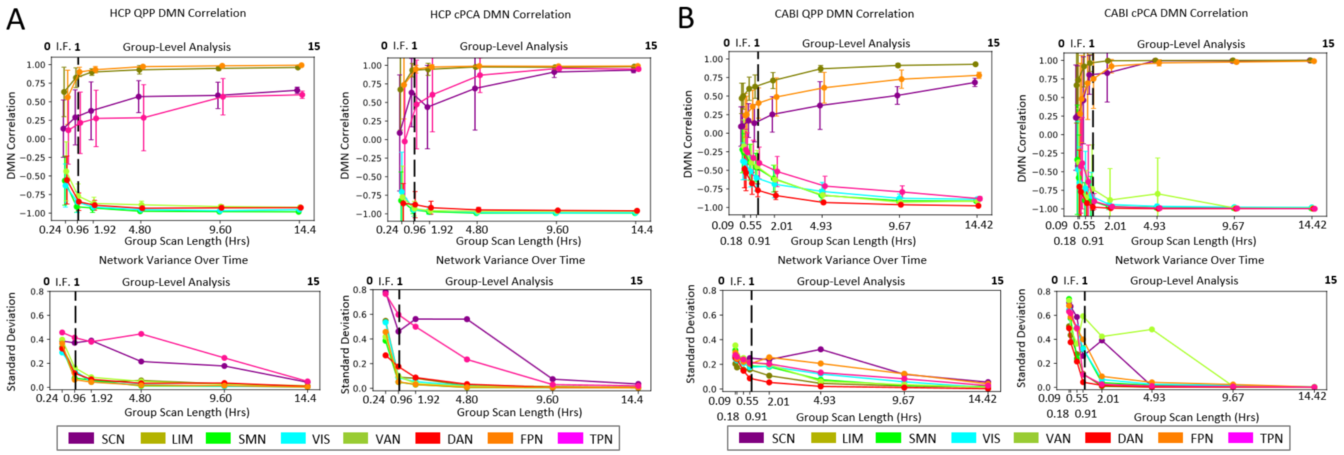


**Figure A1:** *Linear Demonstration of DMN Correlation and Variability Over Scan Length for QPP and cPCA Analyses*. This is the same as Figure **??** with a Linear Scaling for the X-Axis. I.F. (Individual Feasibility) is harder to discern. (A, B) DMN Correlation and Network Variability Over Time. The relationship between the DMN correlation and scan length is assessed for both QPPs and cPCA across two datasets: (A) HCP and (B) CABI Rest. Independent subject groups are used to evaluate variability, with the top row displaying DMN correlation over time and the bottom row showing the standard deviation of network variance over time. The black dashed line marks the methodological transition from individual feasibility to group-level analysis.

#### 1.5 (Objective 1) Scan Length Processing Time for QPPs and cPCA

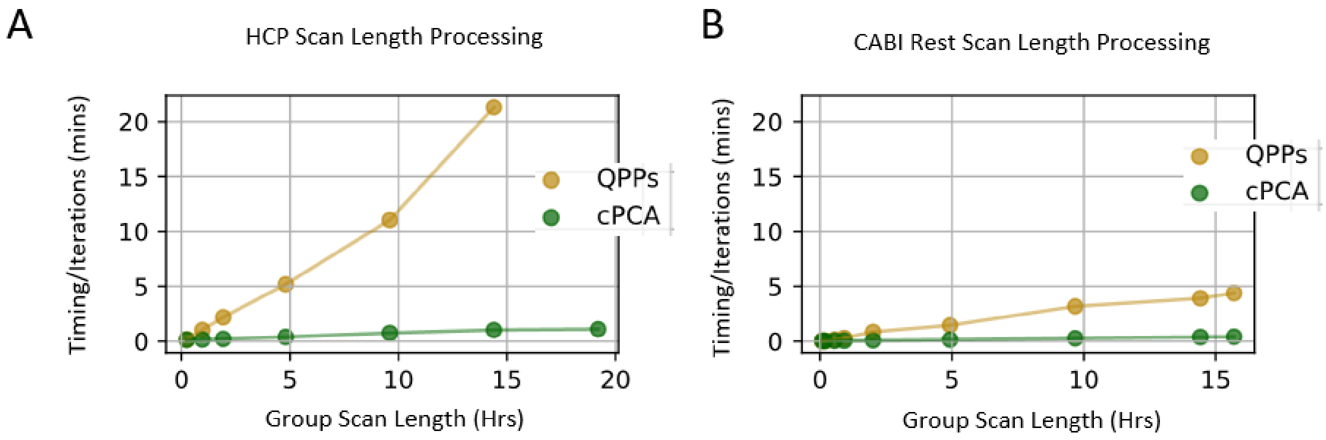


**Figure A2:** *Scan Length Processing Time for QPPs and cPCA* Processing time per iteration is shown for both (A) HCP and (B) CABI Rest datasets. QPPs exhibit linear complexity growth as scan length increases, whereas cPCA follows a quadratic growth pattern. However, due to the relatively low number of timepoints in this analysis, cPCA remains computationally efficient despite its theoretical complexity.

#### 1.6 (Objective 1) Result from DMN Correlation

| Index | Dataset | TR (s) | Algorithm | Group Scan Length (hrs) | Iterations | Avg. Time Per Iter (min) | Total Proc. Time (hrs) |
|---|---|---|---|---|---|---|---|
| 1 | HCP | 0.72 | fastQPP | 0.24 | 80 | 14.17 | 18.89 |
| 2 | HCP | 0.72 | fastQPP | 0.96 | 30 | 31.10 | 15.55 |
| 3 | HCP | 0.72 | fastQPP | 1.92 | 20 | 43.80 | 14.60 |
| 4 | HCP | 0.72 | fastQPP | 4.8 | 15 | 77.82 | 19.45 |
| 5 | HCP | 0.72 | fastQPP | 9.6 | 10 | 110.47 | 18.41 |
| 6 | HCP | 0.72 | fastQPP | 14.4 | 5 | 106.60 | 8.88 |
| 7 | HCP | 0.72 | fastQPP | 19.2 | 1 | 0.00 | 0.00 |
| 8 | HCP | 0.72 | cPCA | 0.24 | 80 | 10.63 | 14.18 |
| 9 | HCP | 0.72 | cPCA | 0.96 | 30 | 4.82 | 2.41 |
| 10 | HCP | 0.72 | cPCA | 1.92 | 20 | 4.37 | 1.46 |
| 11 | HCP | 0.72 | cPCA | 4.8 | 15 | 5.95 | 1.49 |
| 12 | HCP | 0.72 | cPCA | 9.6 | 10 | 7.53 | 1.26 |
| 13 | HCP | 0.72 | cPCA | 14.4 | 5 | 5.13 | 0.43 |
| 14 | HCP | 0.72 | cPCA | 19.2 | 1 | 1.10 | 0.02 |
| 15 | CABI | 2.25 | *robustQPP | 0.091 | 172 | 6.77 | 19.40 |
| 16 | CABI | 2.25 | fastQPP | 0.183 | 80 | 1.88 | 2.51 |
| 17 | CABI | 2.25 | fastQPP | 0.548 | 80 | 11.95 | 15.93 |
| 18 | CABI | 2.25 | fastQPP | 0.913 | 30 | 8.80 | 4.40 |
| 19 | CABI | 2.25 | fastQPP | 2.008 | 20 | 17.22 | 5.74 |
| 20 | CABI | 2.25 | fastQPP | 4.928 | 15 | 21.83 | 5.46 |
| 21 | CABI | 2.25 | fastQPP | 9.673 | 10 | 31.83 | 5.31 |
| 22 | CABI | 2.25 | fastQPP | 14.418 | 5 | 19.58 | 1.63 |
| 23 | CABI | 2.25 | fastQPP | 15.695 | 1 | 4.38 | 0.07 |
| 24 | CABI | 2.25 | cPCA | 0.091 | 172 | 5.92 | 16.96 |
| 25 | CABI | 2.25 | cPCA | 0.183 | 80 | 2.93 | 3.91 |
| 26 | CABI | 2.25 | cPCA | 0.548 | 80 | 3.73 | 4.98 |
| 27 | CABI | 2.25 | cPCA | 0.913 | 30 | 1.73 | 0.87 |
| 28 | CABI | 2.25 | cPCA | 2.008 | 20 | 1.68 | 0.56 |
| 29 | CABI | 2.25 | cPCA | 4.928 | 15 | 2.23 | 0.56 |
| 30 | CABI | 2.25 | cPCA | 9.673 | 10 | 2.73 | 0.46 |
| 31 | CABI | 2.25 | cPCA | 14.418 | 5 | 1.93 | 0.16 |
| 32 | CABI | 2.25 | cPCA | 15.695 | 1 | 0.42 | 0.01 |

**Table A1:** Processing times for fastQPP, robustQPP, and cPCA across different datasets and scan conditions. Group Scan Length refers to the total data duration for each run. Iterations indicate how many times each algorithm was executed per condition. Avg. Time per Iteration measures the computational cost per run. Total Processing Time represents the cumulative computational time for each condition. The (*) indicates the robustQPP algorithm run on these groups since the fastQPP algorithm did not coverage with limited scan time.

#### 1.7 (Objective 1) Complexity Analysis of QPPs and cPCA

##### 1.7.1 Quasi-Periodic Patterns (QPPs)

The computational complexity of QPP detection primarily arises from temporal correlation computations across sliding windows of length $WL$:

1. **Sliding Window Computation**: Segmenting the time series into overlapping windows:

$$O(N \cdot T \cdot WL) \tag{1}$$

2. **Template Matching**: Cross-correlating each windowed segment:

$$O(N \cdot WL \cdot T) \tag{2}$$

3. **Iterative Refinement**: Iterating $K$ times:

$$O(K \cdot N \cdot T \cdot WL) \tag{3}$$

Since complexity scales linearly with $N$ and $WL$, larger window lengths significantly increase computational cost. For practical cases where $WL = 24$:

$$O(256 \times T \times 24) \tag{4}$$

##### 1.7.2 Complex Principal Component Analysis (cPCA)

cPCA performs dimensionality reduction on a complex correlation matrix derived from Hilbert-transformed BOLD signals. The key steps are:

1. **Hilbert Transform** (FFT-based):

$$O(NT \log T) \tag{5}$$

2. **Complex Correlation Matrix Computation**:

$$O(N^2 T) \tag{6}$$

3. **Singular Value Decomposition (SVD)**:

$$O(N^3) \tag{7}$$

4. **Retaining Top $n$ Components**:

$$O(N^2) \tag{8}$$

With iterative convergence over $K$ steps, worst-case complexity is:

$$O(KN^2T) \tag{9}$$

For neuroimaging applications ($N = 256$), efficient SVD implementations keep computation feasible, effectively reducing complexity to:

$$O(N^2T) \tag{10}$$

##### 1.7.3 Single Scan Complexity

For CABI ($T = 146$, $N = 256$, $WL = 24$):

$$O(256 \times 146 \times 24) = O(8.98 \times 10^5) \tag{11}$$

For HCP ($T = 1200$, $WL = 24$):

$$O(256 \times 1200 \times 24) = O(7.37 \times 10^6) \tag{12}$$

For cPCA:

$$O(256^2 \times 146) = O(9.58 \times 10^6) \tag{13}$$

$$O(256^2 \times 1200) = O(7.91 \times 10^7) \tag{14}$$

##### 1.7.4 Full Session Complexity

For CABI ($T = 292$, $WL = 24$):

$$O(256 \times 292 \times 24) = O(1.79 \times 10^6) \tag{15}$$

$$O(256^2 \times 292) = O(1.92 \times 10^7) \tag{16}$$

For HCP ($T = 4800$, $WL = 24$):

$$O(256 \times 4800 \times 24) = O(3.00 \times 10^7) \tag{17}$$

$$O(256^2 \times 4800) = O(3.16 \times 10^8) \tag{18}$$

##### 1.7.5 Worst-Case Group-Level Complexity

For 14-hour datasets ($WL = 24$):

**CABI ($T = 25,112$):**

$$O(256 \times 25112 \times 24) = O(2.52 \times 10^8) \tag{19}$$

$$O(256^2 \times 25112) = O(1.65 \times 10^9) \tag{20}$$

**HCP ($T = 96,000$):**

$$O(256 \times 96000 \times 24) = O(9.65 \times 10^8) \tag{21}$$

$$O(256^2 \times 96000) = O(6.32 \times 10^9) \tag{22}$$

###### 1.7.6 Comparative Complexity Analysis

| Condition | QPP Complexity | cPCA Complexity |
|---|---|---|
| Theoretical Worst Case | $O(K \cdot N \cdot T \cdot WL)$ | $O(KN^2T)$ |
| Single Scan (CABI, $T = 146$) | $O(8.98 \times 10^5)$ | $O(9.58 \times 10^6)$ |
| Single Scan (HCP, $T = 1200$) | $O(7.37 \times 10^6)$ | $O(7.91 \times 10^7)$ |
| Full Session (CABI, $T = 292$) | $O(1.79 \times 10^6)$ | $O(1.92 \times 10^7)$ |
| Full Session (HCP, $T = 4800$) | $O(3.00 \times 10^7)$ | $O(3.16 \times 10^8)$ |
| Group-Level CABI (14 hrs, $T = 25112$) | $O(2.52 \times 10^8)$ | $O(1.65 \times 10^9)$ |
| Group-Level HCP (14 hrs, $T = 96000$) | $O(9.65 \times 10^8)$ | $O(6.32 \times 10^9)$ |

**Table A2:** Computational complexity comparison of QPP and cPCA.

###### 1.7.7 Key Takeaways

QPP scales linearly with $N$, making it highly sensitive to window length ($WL$), whereas cPCA has a quadratic dependence on $N$, though optimized matrix factorization ensures its computational feasibility. For individual scans, such as $T = 146$ for CABI and $T = 1200$ for HCP, both QPP and cPCA remain viable. In full-session scans, where $T = 292$ (2 sessions max per subject from study) for CABI and $T = 4800$ (4 sessions max per subject from study) for HCP, QPP and cPCA exhibit comparable efficiency. However, in worst-case group-level analyses involving 14-hour datasets, with $T = 25,112$ for CABI and $T = 96,000$ for HCP, theoretically cPCA becomes significantly more computationally demanding. Despite this, parallelized and optimized SVD implementations, such as Facebook's fast randomized SVD (fbpca) (Halko et al., 2011), keep computation within practical limits for neuroimaging applications. These methods leverage randomized projections to approximate dominant singular vectors efficiently, reducing complexity compared to classical SVD while maintaining high accuracy.

#### 1.8 (Objective 2) Demonstration of Resampled QPP Templates

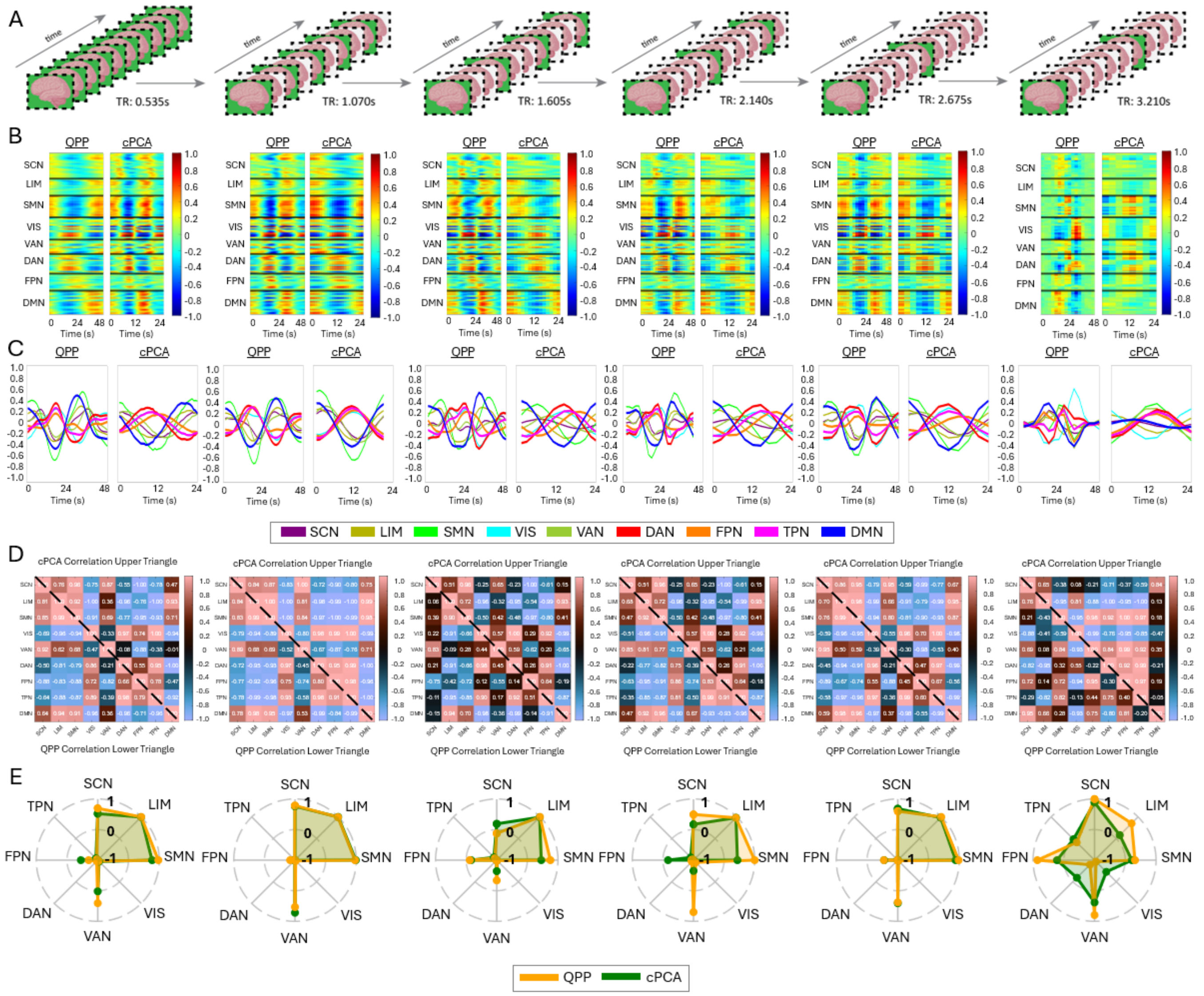


**Figure A3:** *Impact of Temporal Resampling on QPP and cPCA Detection in Videogamer Data.* (A) Illustration of six groups of resampled fMRI data from the Videogamer dataset, demonstrating the effect of increasing TR. TR values range from the native acquisition rate to a degraded TR of 3.210s, allowing for systematic assessment of temporal resolution effects in a dataset with task-related engagement. (B-C) QPP and cPCA template waveforms extracted from each resampled dataset, plotted across the (B) 246 Brainnetome ROIs and (C) eight canonical Yeo networks (SCN, LIM, SMN, VIS, VAN, DAN, FPN, TPN, DMN). The overall waveform structure remains consistent across resampling conditions, though subtle variations emerge with increased TR. (D) Correlation matrices comparing QPP-derived and cPCA-derived templates, with the lower triangular matrix representing QPP correlations and the upper triangular matrix representing cPCA correlations. (E) Polar plots depicting the correlation between the default mode network (DMN) and all other Yeo networks. QPP-derived (yellow) and cPCA-derived (green) correlations are displayed for comparison, showing that the network dynamics between the DMN and the other networks remains preserved even at a resampled TR of 3.210s. These findings highlight that despite task-related modulations, both QPP and cPCA reliably capture large-scale spatiotemporal dynamics, reinforcing their feasibility for group-level analyses even with lower temporal resolution.

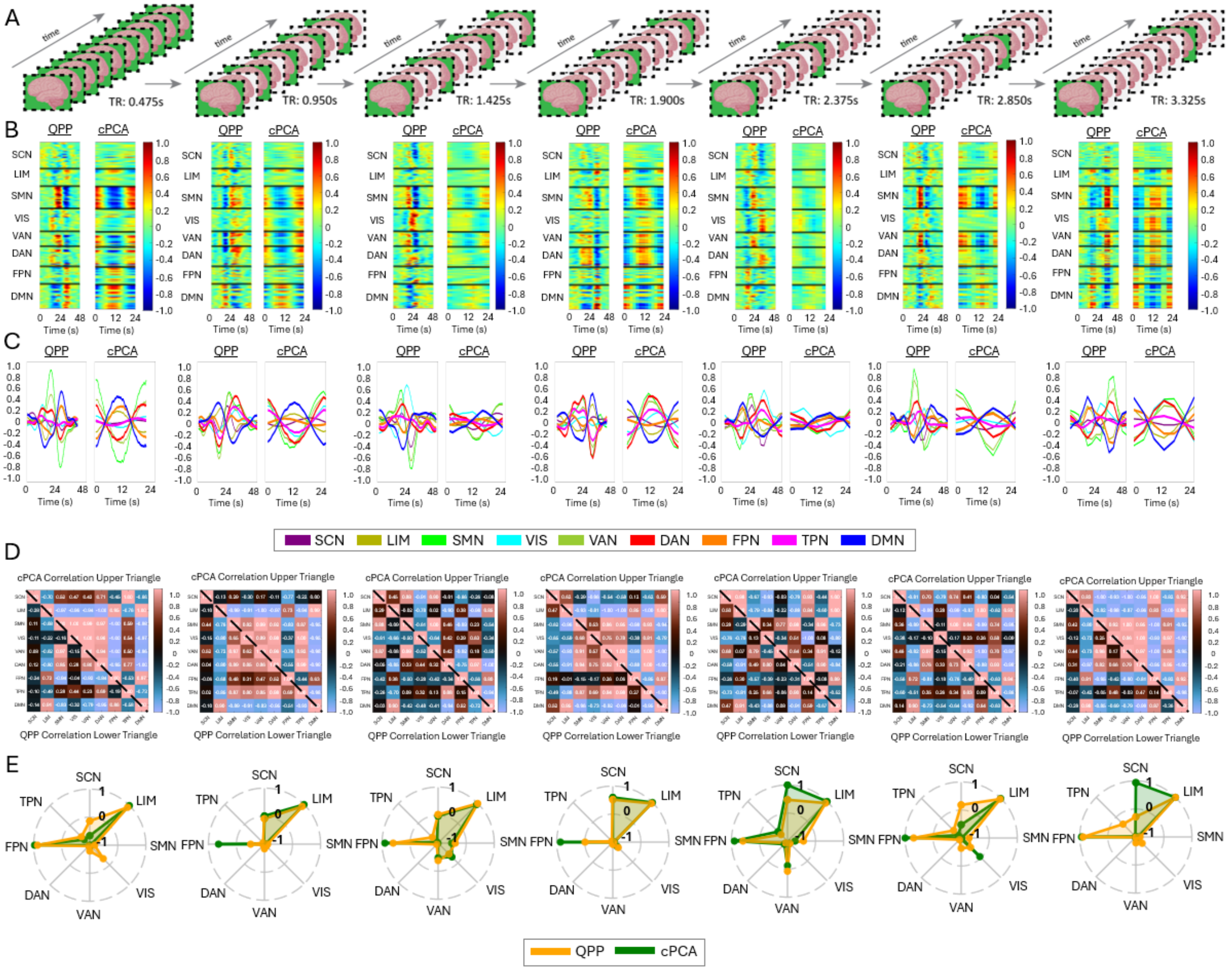


**Figure A4:** *Impact of Temporal Resampling on QPP and cPCA Detection in Musician Data.* (A) Illustration of seven groups of resampled fMRI data from the Musician dataset, demonstrating the effect of increasing TR. TR values range from the native acquisition rate to a degraded TR of 3.325s, allowing for systematic assessment of temporal resolution effects in a shorter scan duration setting. (B-C) QPP and cPCA template waveforms extracted from each resampled dataset, plotted across the (B) 246 Brainnetome ROIs and (C) eight canonical Yeo networks (SCN, LMN, SMN, VIS, VAN, DAN, FPN, TPN, DMN). The overall waveform structure remains consistent across resampling conditions, though subtle variations emerge with increased TR. (D) Correlation matrices comparing QPP-derived and cPCA-derived templates, with the lower triangular matrix representing QPP correlations and the upper triangular matrix representing cPCA correlations. (E) Polar plots depicting the correlation between the default mode network (DMN) and all other Yeo networks. QPP-derived (yellow) and cPCA-derived (green) correlations are displayed for comparison, showing that the network dynamics between the DMN and the other networks remains preserved even at a resampled TR of 3.325s. These findings highlight that despite shorter scan durations, both QPP and cPCA reliably capture large-scale spatiotemporal dynamics, reinforcing their feasibility for individual-level analyses at lower TRs.

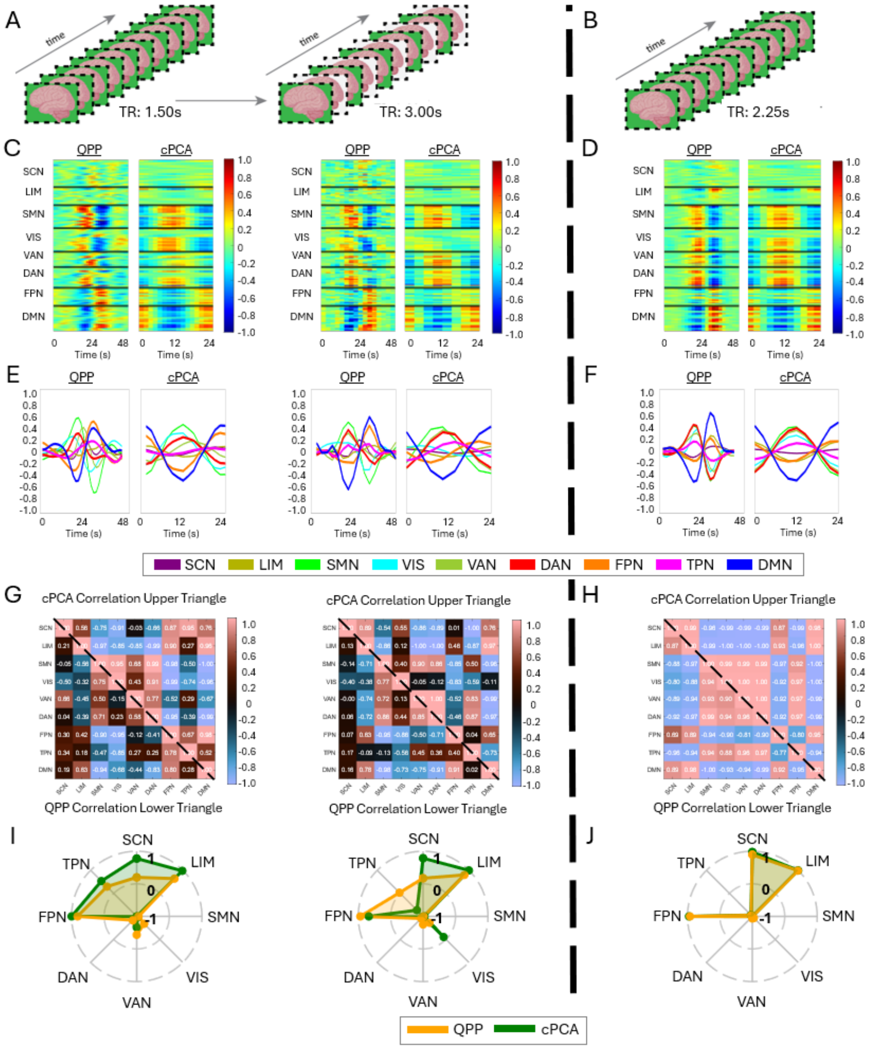


**Figure A5:** *Impact of Temporal Resampling on QPP and cPCA Detection in Meditation and CABI Data.* (A,C,E,G,I) Meditators. (B,D,F,H,J) CABI Rest Data. (A,B) Illustration of two groups and one group of resampled fMRI data from the Meditation and CABI datasets, demonstrating the effect of increasing TR. TR values range from the native acquisition rate to a degraded TR of 3.00s, allowing for systematic assessment of temporal resolution effects in both structured meditation paradigms and naturalistic resting-state conditions. (C-F) QPP and cPCA template waveforms extracted from each resampled dataset, plotted across the (C,D) 246 Brainnetome ROIs and (E,F) eight canonical Yeo networks (SCN, LMN, SMN, VIS, VAN, DAN, FPN, TPN, DMN). The overall waveform structure remains consistent across resampling conditions. (G,H) Correlation matrices comparing QPP-derived and cPCA-derived templates, with the lower triangular matrix representing QPP correlations and the upper triangular matrix representing cPCA correlations. (I,J) Polar plots depicting the correlation between the default mode network (DMN) and all other Yeo networks. QPP-derived (yellow) and cPCA-derived (green) correlations are displayed for comparison, showing that the network dynamics between the DMN and the other networks remains preserved even at a resampled TR of 3.00s for the meditation group. These findings highlight that both QPP and cPCA reliably capture large-scale spatiotemporal dynamics, reinforcing their feasibility for analyzing lower TR-sampled data in varied mental states.

#### 1.9 (Objective 2) Values for QPP and cPCA Significant Test

| Dataset | t-statistic | p-value |
|---|---|---|
| HCP | -0.197278 | 0.844129 |
| Videogamers | 0.237193 | 0.813024 |
| Musicians | -0.591847 | 0.555189 |
| Meditators | -0.331465 | 0.742698 |
| CABI Rest | 0.006748 | 0.994711 |

**Table A3:** Statistical comparison between QPPs and cPCA across the five datasets. The t-statistic and p-values indicate whether significant differences exist between algorithms. The results suggest fidelity between the algorithms, as none of the comparisons yield statistically significant differences across intrinsic network dynamics.

#### 1.10 (Objective 3) Modified Jaccard Index Demonstration

| Band | TMX1 (Maxima) | TMX2 (Minima) |
|---|---|---|
| Infraslow+ | 467 | 441 |
| Infraslow | 506 | 472 |
| Slow5 | 384 | 365 |
| Slow4 | 581 | 571 |

| Comparison | TMX1 (Maxima) | TMX2 (Minima) |
|---|---|---|
| Infraslow+ and Infraslow | 0.4424 | 0.4447 |
| Infraslow+ and Slow5 | **0.1969*** | **0.1950*** |
| Infraslow+ and Slow4 | 0.3311 | 0.3241 |
| Infraslow and Slow5 | **0.1059*** | **0.0944*** |
| Infraslow and Slow4 | 0.3554 | 0.3574 |
| Slow5 and Slow4 | **0.2042*** | **0.1841*** |

**Table A4:** Comparison of TMX1 (Maxima) and TMX2 (Minima) instances for each frequency band (left) and the Modified Jaccard Index for Maxima (TMX1) and Minima (TMX2) with Cushion ($\pm 16$ timepoints) (right). Statistical testing revealed no significant difference between TMX1 and TMX2, as assessed by the Wilcoxon signed-rank test ($p = 0.31250$). However, the Kruskal-Wallis test indicated a significant difference between Slow-5 and other frequency bands ($p = 0.00216$), suggesting that Slow-5 exhibits distinct spatiotemporal properties compared to the other bands. The **bold*** values indicate statistically significant differences between groups using pair-wise comparison. All groups containing Slow-5 were not cyclically aligned with the other frequencies.

#### 1.11 (Objective 3) Instances and Characteristics of QPPs throughout Data

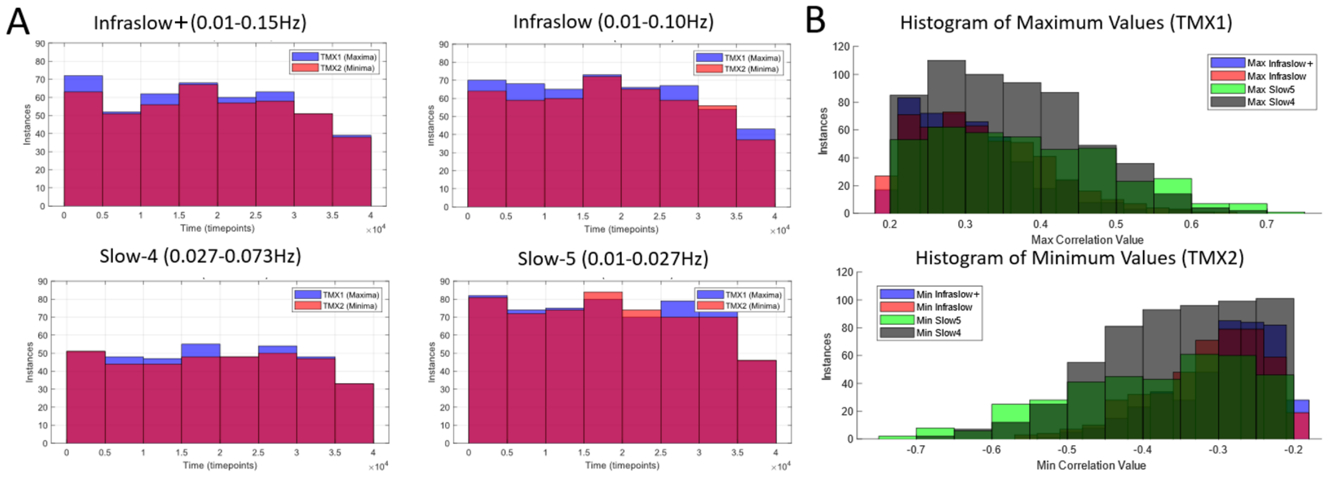


**Figure A6:** *Distribution of QPP Detection Across Frequency Bands in the Concatenated Group.* (A) Temporal distribution of detected QPP maxima (TMX1, blue) and minima (TMX2, red) across all four frequency bands. The relatively uniform distribution across the time axis suggests that QPP patterns are consistently detected throughout the scan, with no substantial clustering in specific time intervals. Moreover, the comparable proportions of TMX1 and TMX2 indicate that network fluctuations are evenly distributed between high and low correlation states across all frequency bands. (B) Histograms of the maximum (top) and minimum (bottom) correlation values for each frequency band. The distributions reveal no major differences in the overall spread or magnitude of maxima and minima across frequency bands, further supporting the observation that QPP detection remains stable regardless of the selected frequency band. These findings suggest that QPPs are robustly identified across different frequency bands and exhibit a consistent balance between high and low network correlation states.

#### 1.12 (Objective 3) Testing Window Length for Slow-5 Frequencies

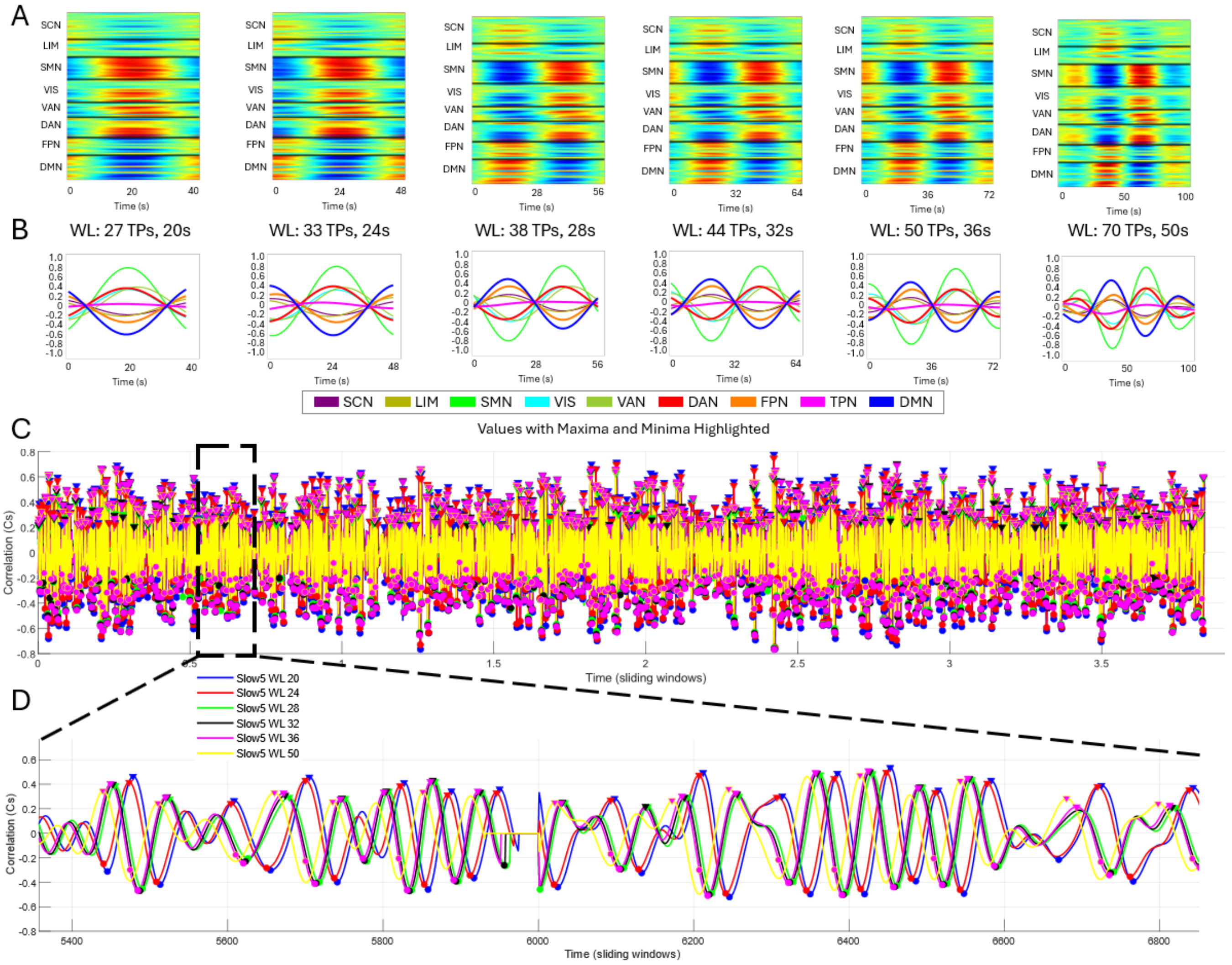


**Figure A7:** *Impact of Window Length on QPP Detection in the Slow-5 Frequency Band of HCP Data.* (A-B) QPP templates extracted using different window lengths (WLs) within the Slow-5 frequency band across the (A) 246 Brainnetome ROIs and (B) eight Yeo networks. The range tested includes window lengths spanning 20–36 seconds, with a focus on evaluating the transition within the typical QPP range (20–30s) as well as longer WLs beyond this range (32s, 36s, and 50s). (C) Group-level QPP detection across the concatenated HCP dataset, highlighting the temporal locations of detected QPPs for different window lengths. Maxima and minima are indicated for each detected QPP, showing how detection patterns vary with changing WL. (D) A zoomed-in segment of the time series, illustrating the correlation structure of Slow-5 QPPs across different window lengths. Notably, around 26 seconds, a shift is observed in network synchronization, transitioning from capturing structured inter-network correlations to detecting spontaneous network state transitions. This suggests that longer window lengths may be identifying a distinct, slower dynamic change in the QPP structure. These findings highlight the potential for future research into the role of window length in detecting network dynamics within the Slow-5 frequency range and its implications for modeling large-scale brain activity.

#### 1.13 (Objective 3) cPCA Frequency Analysis

| Exp. Var. | PCA1 | PCA2 | PCA3 | PCA4 | PCA5 | PCA6 | PCA7 | PCA8 | PCA9 | PCA10 |
|---|---|---|---|---|---|---|---|---|---|---|
| Infraslow+ | 0.228 | 0.122 | 0.102 | 0.080 | 0.063 | 0.059 | 0.045 | 0.040 | 0.037 | 0.028 |
| Infraslow | 0.243 | 0.133 | 0.107 | 0.085 | 0.066 | 0.063 | 0.047 | 0.042 | 0.039 | 0.030 |
| Slow5 | 0.266 | 0.166 | 0.116 | 0.096 | 0.080 | 0.072 | 0.054 | 0.044 | 0.040 | 0.036 |
| Slow4 | 0.254 | 0.131 | 0.110 | 0.084 | 0.066 | 0.062 | 0.048 | 0.041 | 0.039 | 0.030 |

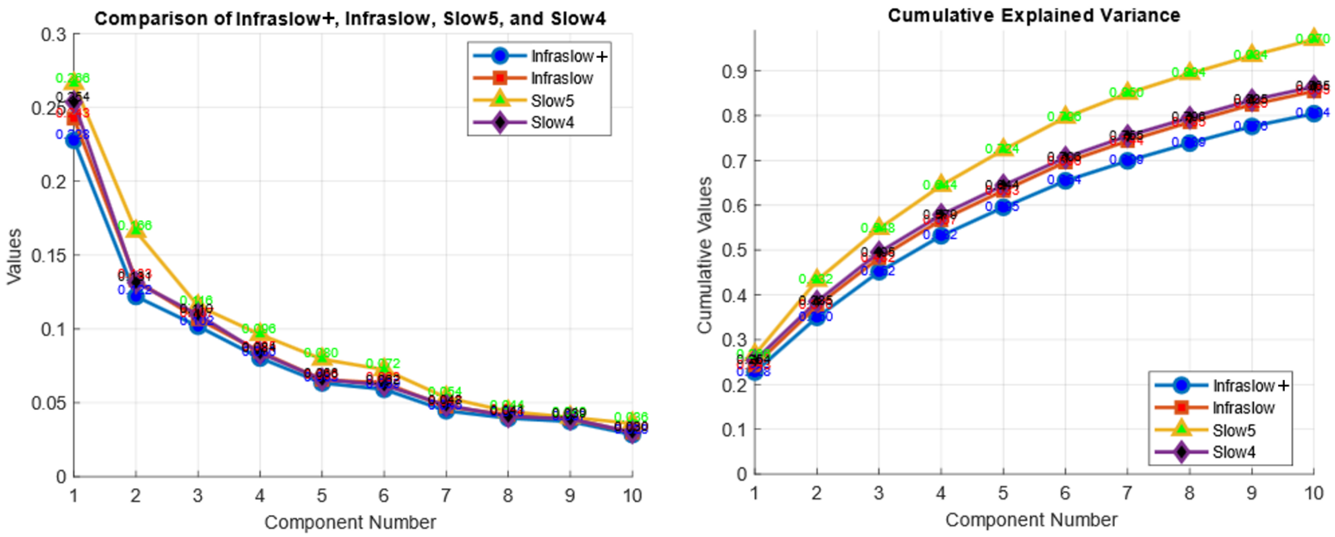


**Figure A8:** Explained variance for the first 10 principal components across different frequency bands (top). Below are the two corresponding figures illustrating the results.
